# A mathematical framework for modelling the metastatic spread of cancer

**DOI:** 10.1101/469536

**Authors:** Linnéa C Franßen, Tommaso Lorenzi, Andrew EF Burgess, Mark AJ Chaplain

**Keywords:** Metastatic spread, Mathematical oncology, Tumour microenvironment, Individual-based model, Multi-grid framework

## Abstract

Cancer is a complex disease that starts with mutations of key genes in one cell or a small group of cells at a primary site in the body. If these cancer cells continue to grow successfully and, at some later stage, invade the surrounding tissue and acquire a vascular network (tumour-induced angiogenesis), they can spread to distant secondary sites in the body. This process, known as *metastatic spread*, is responsible for around 90% of deaths from cancer and is one of the so-called *hallmarks of cancer*.

To shed light on the metastatic process, we present a mathematical modelling framework that captures for the first time the interconnected processes of invasion and metastatic spread of individual cancer cells in a spatially explicit manner — a multi-grid, hybrid, individual-based approach. This framework accounts for the spatio-temporal evolution of mesenchymal- and epithelial-like cancer cells, as well as MT1-MMP and MMP-2 dynamics, and interactions with the extracellular matrix.

Using computational simulations, we demonstrate that our model captures all the key steps of the invasion-metastasis cascade, i.e. invasion by both heterogeneous cancer cell clusters and by single mesenchymal-like cancer cells; intravasation of these clusters and single cells both via active mechanisms mediated by matrix degrading enzymes (MDEs) and via passive shedding; circulation of cancer cell clusters and single cancer cells in the vasculature with the associated risk of cell death and disaggregation of clusters; extravasation of clusters and single cells; and metastatic growth at distant secondary sites in the body. By faithfully reproducing experimental results, our simulations support the evidence-based hypothesis that the membrane-bound MT1-MMP is the main driver of invasive spread rather than diffusible MDEs like MMP-2.

## 1 Introduction

Most solid tumours start with mutations of key genes either in one or in a small group of the more than 10^14^ healthy cells in the human body. Abnormally rapid cell proliferation is one of the most notable results of these acquired cancerous mutations, which can lead to the formation of a small nodule of cancer cells. Over time, this nodule can expand, while acquiring increasingly aggressive mutations, into a full avascular tumour with a diameter of up to approximately 0.1–0.2 cm (Folkman, 1990), limited by the diffusion of nutrients (e.g. oxygen). For successful growth beyond this size, the cancer cells start recruiting new blood vessels by secreting chemicals — collectively known as tumour angiogenic factors (TAFs) (Folkman and Klagsbrun, 1987). This neovascularisation process is called (*tumour-induced) angiogenesis*. The newly established vasculature serves the tumour’s increased metabolic needs by transporting the required nutrients. The newly formed blood vessels additionally benefit the tumour in the subsequent *vascular growth phase*, when the cancer cells become invasive so that gradients in nutrients, oxygen and extracellular matrix (ECM) drive cancer cells away from the primary tumour mass. In the event that cancer cells successfully intravasate into the newly grown blood vessels *and* survive in the vessel environment (where they are exposed to risks such as attacks by the immune system and shear stress in the blood flow), they can extravasate and relocate at distant sites in the body. At these new sites, nutrients and space are less of a limiting factor to growth. The described sequence of steps of successful relocation of cancer cells from a primary location to a secondary location is known as *metastatic spread*. It can lead to the formation of secondary tumours, called *metastases*, at sites in the body away from the primary tumour. In the first instance, however, the successfully extravasated cancer cells occur either as single *disseminated tumour cells* (DTCs) or as small clusters of cancer cells, called *micrometastases*. These DTCs and micrometastases may remain dormant but have the potential to proliferate into vascularised *metastases* at the metastatic sites at some later point in time. The full process we described here, which is shown schematically in Figure 1, is also known as the *invasion-metastasis cascade* (Fidler, 2003; Talmadge and Fidler, 2010).

**Fig. 1:**
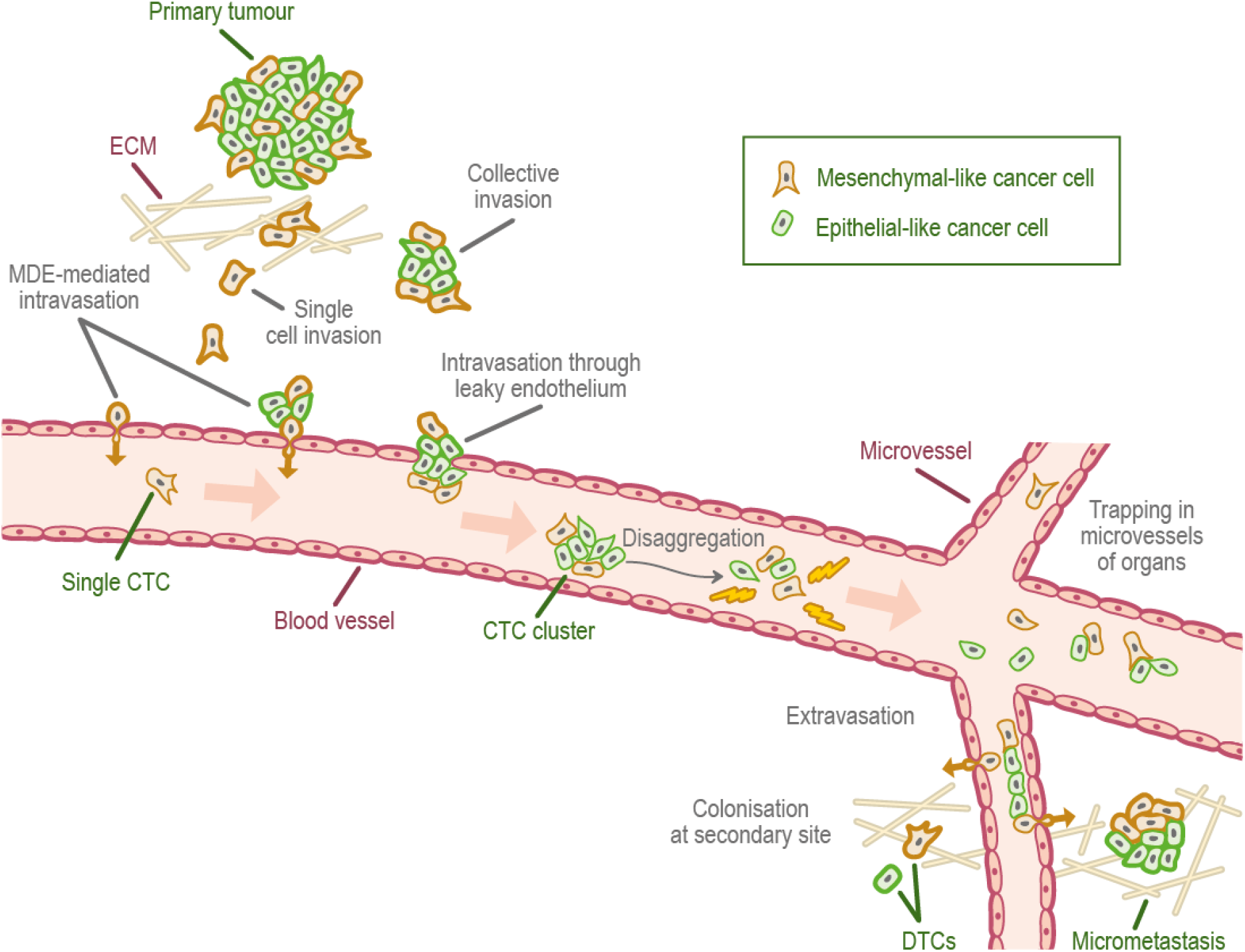
Schematic overview of the invasion-metastasis cascade. Single mesenchymal-like cancer cells and heterogeneous clusters of mesenchymal- and epithelial-like cancer cells break free from the primary tumour and invade the surrounding tissue (top left). They can intravasate via active MDE-mediated and passive mechanisms (upper left, epithelium of vessel). Once in the vasculature, CTC clusters may disaggregate (centre). Extravasation to various secondary sites in the body is rare. However, successful colonisation there can result in either DTCs or micrometastases (bottom right), which have the potential to develop into full-blown metastases.

Expanding and deepening our understanding of the invasion-metastasis cascade is of vital importance. Only approximately 10% of cancer-related deaths are caused by primary tumours alone that, for example, have grown to a size at which they affect organ function by exerting physical pressure. Although this alone is a incentive to model cancer growth, the other 90% of cancer-related deaths arise due to metastatic spread and metastases growing at distant sites away from the primary tumour (Hanahan and Weinberg, 2000; Gupta and Massagué, 2006). Many localised primary tumours can be treated successfully, e.g. by resection or chemotherapy, but once cancer cells have begun to spread throughout the body, it becomes increasingly difficult to treat a patient successfully and prognosis is very poor.

The invasion-metastasis cascade is a complex biological process and many questions about its details remain unanswered to date. Mathematical modelling can therefore be a useful tool to capture and unravel this complexity, and to thereby gain a better understanding of the invasion-metastasis cascade. Ultimately, predictive modelling may help to advance treatment success through personalised medicine.

While spatial mathematical models of the cancer invasion process alone as well as non-spatial models of metastatic spread exist, these two processes, both of which are spatial in nature, have, to our knowledge, not been combined into a unified mathematical modelling framework previously. In this paper, we hence propose a first such framework to model cancer cell invasion and metastatic spread in a explicitly spatial manner. We consider cancer cells to be of two phenotypes — the more motile *mesenchymal-like* and the more proliferative *epithelial-like* cancer cells. Adopting a multi-grid, hybrid, individual-based approach, our mathematical model is capable of capturing all the key steps of the invasion-metastasis cascade, i.e. invasion by both heterogeneous cancer cell clusters and by single mesenchymal-like cancer cells; intravasation of these clusters and single cells both via active mechanisms, which are mediated by matrix degrading enzymes (MDEs), and via passive shedding; circulation of cancer cell clusters and single cancer cells in the vasculature with the associated disaggregation of clusters and risk of cell death; extravasation of clusters and single cells; and metastatic growth at distant sites in the body. We verify the biological accuracy of our model by varying key model parameters and checking results against biological findings. We specifically find that the co-presence of epithelial-like and mesenchymal-like cancer cells enhances local tissue invasion and metastatic spread. Furthermore, by reproducing experimental results, our simulations support the evidence-based hypothesis that the membrane-bound MT1-MMP is the main driver of invasive spread rather than diffusible MDEs like MMP-2, in particular when we switch from diffusion-dominated to haptotaxis-dominated cancer cell invasion.

The remainder of the paper is organised as follows. In Section 2, we summarise the key steps of the invasion-metastasis cascade and the roles of different cancer cell phenotypes in more detail. In Section 3, we introduce our general mathematical modelling framework of metastatic spread. As part of introducing this new modelling framework, we give an overview of previous models of both cancer invasion and metastasis at the beginning of this section. In Section 4, we explain how we set up the computational simulations and how we calibrated the model. In Section 5, we present the simulation results. Finally, in Section 6, we discuss the biological implications of our results as well as future work.

## 2 Biological background

In this section, we will describe the main steps of the *invasion-metastasis cascade* (*cf*. Hanahan and Weinberg (2000, 2011)) that can enable a small primary nodule of cancer cells to spread to distant sites throughout the body and then colonise at these secondary sites.

## Local cancer cell invasion

When the cancer cells are invading the tissue surrounding the primary tumour, they have to overcome structural obstacles. The cells need to make their way through the fairly rigid extracellular matrix (ECM), which mainly consists of various tissue-bound macromolecules such as structure-providing collagens (mainly of type-I), as well as of fibronectin, vitronectin and laminin, which influence cancer cell spreading, motility and cell-adhesion. Often, the cancer cells additionally have to penetrate the even more rigid basal laminae of blood vessels and potentially of the primary sites they originate from.

The two main mechanisms used by cancer cells to overcome these hurdles, which have been discussed in detail by, for example, Friedl and Wolf (2003), are *protease-dependent* and *protease-independent* invasion. Protease-dependent invasion earns its name from collagen-cleaving proteinases, and more specifically MDEs, which are secreted by some cancer cell types. The cleaving of collagen allows all types of cancer cells to subsequently invade along the paths created. Protease-independent invasion relies on cancer cells changing from a mesenchymal-like to an amoeboid-like shape — a process called *mesenchymal-amoeboid transition*. This increases the morphological plasticity of the cells and enables them to squeeze through the collagen-like pores, rather than solely relying on ECM degradation.

It has been shown by Sabeh et al (2009) that cancer cells cannot migrate unless the proteinases have cleared the collagen prevalent in normal tissue of its covalent cross-links, and that protease-dependent invasion on its own is a sufficient invasion mechanism. Therefore, we focus only on the mechanism of protease-dependent invasion throughout the rest of this paper.

We distinguish between two cancer cell phenotypes — epithelial-like cancer cells and mesenchymal-like cancer cells. These cancer cell types arise due to an observed tradeoff between a cell’s invasiveness and its ability to proliferate — this is also known as the *go-or-grow dichotomy* (Giese et al, 1996). The mesenchymal-like cancer cells resemble mobile cells in embryo development and are therefore more motile. The mesenchymal-like cancer cells can also invade and intravasate as single cells. This is due to their loss of cell-cell adhesion as well as their expression of MDEs, such as the membrane-bound MT1-MMP and the diffusive MMP-2. Epithelial-like cancer cells, on the other hand, cannot invade effectively without the coexistence of MDE-secreting mesenchymal-like cancer cells. This is because the cancer cells with an epithelial-like phenotype do not express MDEs. They are also comparatively less motile. However, the epithelial-like cell type is more proliferative and its role should not to be ignored in the invasion-metastasis cascade. Also, mesenchymal-like cancer cells have been suggested to be able to develop from epithelial-like cancer cells via a process termed the *epithelial-mesenchymal transition* (EMT) (Kalluri and Weinberg, 2009), which is shown schematically in Figure 2. The reverse process, the *mesenchymal-epithelial transition* (MET), has additionally been suggested to be involved in metastatic spread, for instance by contributing to the colonisation of DTCs at secondary sites (Gunasinghe et al, 2012).

**Fig. 2:**
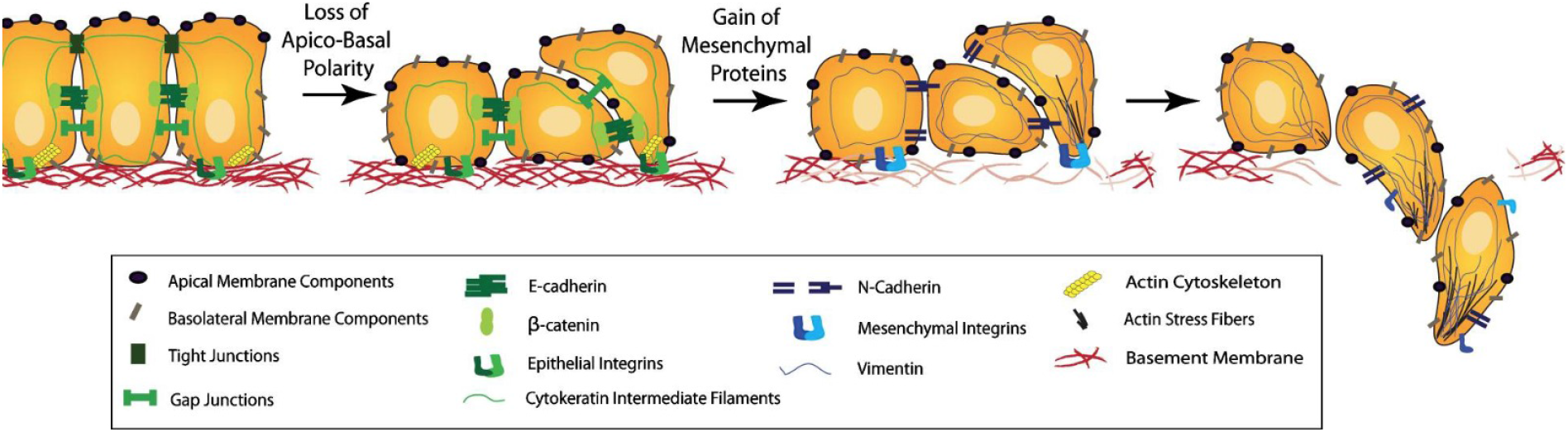
Schematic representation of the EMT. As an outcome of EMT, the cell-cell adhesion between formerly epithelial-like cancer cells is reduced, while the cancer cells express more cell-matrix adhesion enhancing molecules like cadherin. This combination of changes enhances invasiveness. Further, cancer cells become more potent at degrading the underlying basement membranes of organs and vessels, as shown towards the right of the figure, as well as the ECM in general. This allows the mesenchymal-like cancer cells to invade into the surrounding stroma. Reproduced from (Micalizzi et al, 2010) with permission from Springer.

Mesenchymal-like and epithelial-like cancer cells have been observed to invade most effectively in a setting where both cancer cell types are present. This gives rise to the hypothesis of a second protease-dependent invasion mechanism in addition to mesenchymal-like cancer cells invading individually. *Collective migration* of cohesive cell cohorts has been shown to be important for the invasion of cancer cells by Friedl et al (2012), amongst others. The theory is that clusters consisting of cancer cells of heterogeneous phenotypes may invade the extracellular matrix together. Figure 1 provides a scheme of the invasion of single mesenchymal-like cells versus collective groups of cells.

### Intravasation

Once suitably mutated cancer cells have managed to invade the tissue far enough to find themselves adjacent to a lymph or blood vessel (whether in the form of oligoclonal clusters derived from the same primary tumour (Aceto et al, 2014) or individually), they can potentially intravasate into the blood system through the basal laminae of blood vessels. While there is experimental evidence suggesting that a subset of cancer cell lines may only be able to access the blood vessels indirectly via prior intravasation into the lymph vessels, the spread to distant sites in the body ultimately happens by dissemination through the blood vessels (Wong and Hynes, 2006; Lambert et al, 2017).

The exact mechanism of intravasation into the vasculature is still unclear. However, the two main proposed biological intravasation modes — *active* versus *passive* intravasation — are likely not mutually exclusive, as suggested for instance by Cavallaro and Christofori (2001), Bockhorn et al (2007) and Jie et al (2017). The *active* intravasation hypothesis postulates that cancer cells crawl towards and into vessels actively with help of MDEs. *Passive* intravasation, on the other hand, implies a more accidental shedding of cancer cells via newly formed, immature vessels, which are fragile and may collapse due to trauma or under the physical pressure caused by rapid tumour expansion.

The above-explained difference between more mesenchymal-like and more epithelial-like cancer cells together with this differentiation between active and passive intravasation gives rise to three entry modes of cancer cells into the vasculature, which are further explained in (Francart et al, 2018):

1. Single MDE-expressing mesenchymal-like cancer cells actively enter the blood vessels and thereafter disseminate as single circulating tumour cells (CTCs).
2. Epithelial-like and mesenchymal-like cancer cells cooperate in the sense that mesenchymal-like cells allow epithelial-like cells to enter the vasculature together with them, or shortly after them. Mesenchymal-like cells express the MDEs required to degrade the vessels’ basal laminae. This allows for co-invasion of the epithelial-like cancer cells in the vicinity. Thus, both mesenchymal-like and epithelial-like cancer cells enter the blood system jointly as a cluster.
3. Any single cancer cells or cancer cell clusters near a ruptured blood vessel intravasate via the passive entry mode.

These entry mechanisms are depicted — left to right — along the upper left blood vessel wall in Figure 1.

### Travel through the vasculature and extravasation

Successful intravasation into the vasculature by no means implies that the respective cancer cells will succeed in metastasising. Cancer cells encounter further obstacles in the bloodstream. In fact, as Figure 3 shows, single CTCs and CTC clusters are exposed to physical stresses and are attacked by natural killer (NK) cells. This causes cancer cell clusters to disaggregate, as shown in the centre of Figure 1, leading to smaller CTC clusters and an increased number of single CTCs. Also, it further leads to a significant decrease in the number of cancer cells that reach the metastatic site from the primary tumour.

**Fig. 3:**
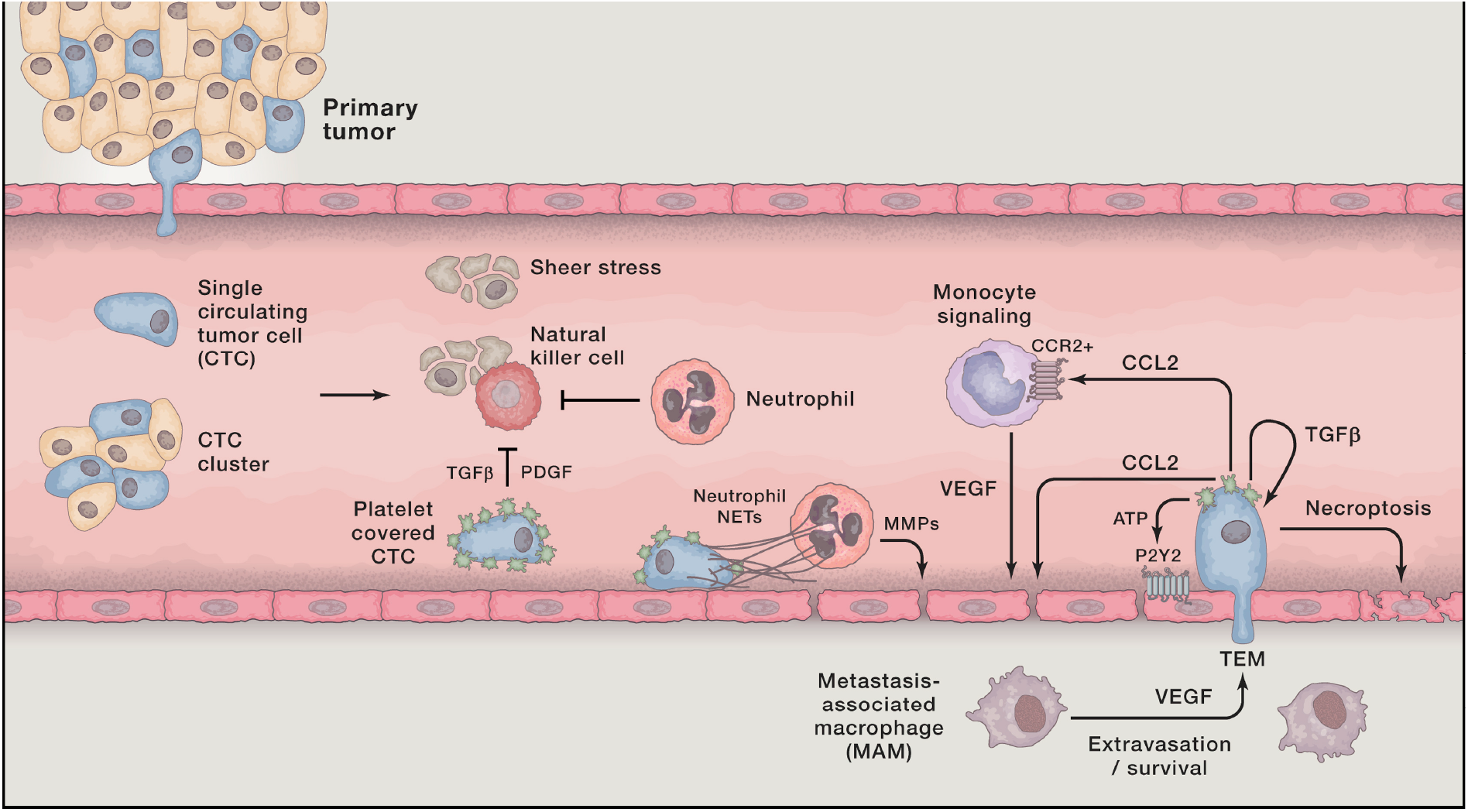
Cancer cells in the blood system. Once single cancer cells or cancer cell clusters have intravasated, a number of mechanisms — both to aid the cancer cells (e.g. platelets covering cell surface, neutrophils that enhance extravasation through NET expression, and MMP secretion) and to destroy them (e.g. physical stresses; attacks by NK cells) — come into action. Reproduced from (Lambert et al, 2017) with permission from Elsevier Inc.

Other cells in the bloodstream assist the cancer cells. Platelets coat the surfaces of the cancer cells, which prevents NK cells from recognising and destroying them. Neutrophils have a similar effect and additionally support metastatic seeding. As depicted in the middle of the lower vessel wall shown in Figure 3, neutrophils can express neutrophil extracellular traps (NETs), which entangle cancer cells. This is suggested to enhance the cancer cells’ survival potential, as well as the probability that they will adhere to endothelial cells and extravasate. Neutrophils also secret various metalloproteases (MMPs) upon arrest, which aid extravasation of the cancer cells by cleaving the vessel wall. Transendothelial migration (TEM) is further provoked by bioactive factors (e.g. VEGF, MMPs and ADAM12), which are secreted by activated platelets and by cancer cells. These factors can act on cancer cells themselves, on monocytes and on endothelial cells. Inflammatory monocytes promote TEM by differentiating into metastasis-associated macrophages that reside in the parenchyma of the potential secondary sites. Finally, it has recently been found by Strilic et al (2016) that cancer cells can induce necroptosis of healthy endothelial cells, as shown on the bottom right of Figure 3, which allows the cancer cells to extravasate without TEM. More in-depth information on the biological background of extravasation can be found in (Lambert et al, 2017).

### Metastatic spread and colonisation

A successfully extravasated single cancer cell or cluster of cancer cells can either contribute to *self-seeding* to an existing metastasis or to the primary tumour, or it can settle as a potential initial seed of a new metastasis (Pantel and Speicher, 2016). However, even if a cancer cell has extravasated successfully into the parenchyma of a potential new metastatic site, success in growing into a full-blown secondary tumour is not guaranteed. In fact, to give a rough idea of the probability that a cancer cell, which has already intravasated successfully, will ultimately develop into a micro- or macrometastasis, we can quote the result of an experimental study by Luzzi et al (1998). The authors studied the proportions of melanoma cells that formed micro- and macrometastases after the cells were injected intra-portally to target mouse livers. They found that 2.04% of the injected cancer cells formed micrometastases after 3 days but that after 13 days only 0.07% of the initially injected cancer cells were still present as micrometastases. Additionally, 0.018%±0.017% of initially injected cancer cells had formed macrometastases after 13 days. These survival probabilities for single CTCs, may, according to Valastyan and Weinberg (2011), even be an overestimation. CTC clusters were described to have between 23 and 50 times the metastatic potential of single CTCs (Aceto et al, 2014). Note, however, that this is only a rough estimate and will depend on other factors such as the particular cancer cell lines and the secondary sites involved, as explained in detail below.

The main mechanisms, by which successfully extravasated cancer cells are prevented from forming malignant secondary tumours, are related to them being maladapted to their new microenvironment, as first suggested by the “seed and soil hypothesis” of Paget (1889). Hence, most cancer cells will be eliminated from the parenchyma at the secondary site. Others will stay at the distant site for periods lasting up to decades but will remain in a dormant state. They fail to proliferate at the secondary site as they are, for example, actively inhibited by the immune system or fail to induce angiogenesis. These — at least transiently — indolent cancer cells can exist either as single DTCs or in the form of micrometastases.

**Fig. 4:**
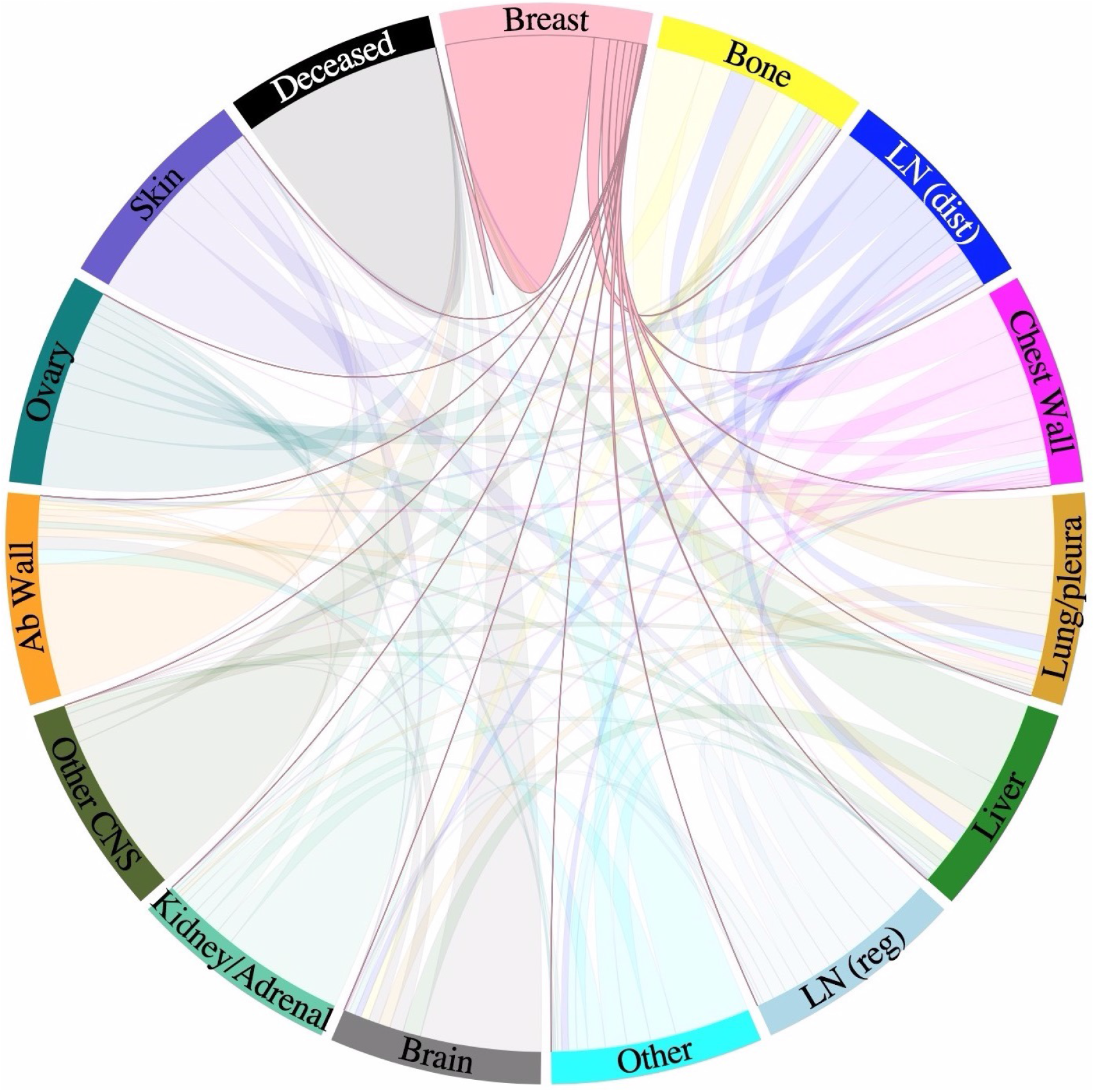
Metastatic progression of breast cancer. Circular chord diagram showing Markov chain network of data on metastatic spread from 4181 breast cancer patients over a 10-year period. Primary breast cancer is located on top with metastatic sites — including bone, lung and brain — ordered clockwise in decreasing order according to transition probability from the primary breast tumour. Chord widths at the ‘breast’ starting location represents one-step transition probabilities between two sites. Further information on the exact data origin and patient criteria can be found at http://kuhn.usc.edu/breast_cancer/. Courtesy of Dr. Jeremy Mason, University of Southern California using the interactive tool published at http://kuhn.usc.edu/forecasting — the corresponding publication is (Newton et al, 2013).

The mechanisms of and the reasons for cancerous spread to specific metastatic sites are largely unknown. However, there are studies that can provide insight into typical patterns of metastatic tumour spread of a certain primary cancer type. To tie in with Paget’s more than 100 year old observation that breast cancer spread does not occur randomly (Paget, 1889), we can, for example, consult data on the metastatic spread of primary breast cancer that were collected from 4181 breast cancer patients (3735 early stage breast cancer patients diagnosed at MD Anderson Cancer Center and 446 breast cancer patients at Memorial Sloan Kettering Cancer Center, who had no detectable metastases upon diagnosis but all developed some eventually) and visualised in interactive graphs by (Kuhn Laboratory, 2017). Figure 4 presents a snapshot of such an interactive figure, which shows typical metastatic spread patterns of a primary breast tumour after 10 years.

## 3 The mathematical modelling framework

In this section, we introduce our general spatial modelling framework of the metastatic spread of cancer. We begin by giving an overview of previous models of cancer invasion and subsequently of metastasis. Throughout, we distinguish models which include only *local* interactions between cancer cells and their environment and those that additionally capture interactions that are *non-local* in space.

Several continuous *local* models formulated in terms of differential equations have been proposed. Following early ordinary differential equation (ODE) models (Gatenby, 1991, 1995a,b), a first one-dimensional spatially explicit model was introduced in the seminal work by Gatenby and Gawlinski (1996). Their paper described spatiotemporal acid-mediated invasion using a system of reaction-diffusion-taxis partial differential equations (PDEs). Perumpanani et al (1996) published a first model of several that considered random motility, haptotaxis and chemotaxis in the context of invasive cancer cells interacting with MDEs, ECM proteins, normal cells and non-invasive cancer cells. The model examined how deeply and at which speed cancer cells invaded the ECM when led by hapto- and chemotactic cues. Ambrosi and Preziosi (2002) proposed a model, in which a tumour spheroid was seen as a growing and deformable porous medium. Using this approach, they introduced a multiphase mechanical framework of tumour growth. A much-cited model focussing on haptotaxis is the continuum PDE model by Anderson et al (2000), which to our knowledge was the first of its kind to extend modelling to a 2D setting. A number of subsequent continuum reaction-diffusion-taxis PDE models focussed on specific MDEs. For instance, the role of urokinase-type plasminogen activator, which is one of the proteolytic enzymes over-expressed in cancer cells, in cancer invasion was studied by Chaplain and Lolas (2005, 2006) using systems of reaction-diffusion-taxis PDEs. Work by Andasari et al (2011) extended this modelling approach. Further, Deakin and Chaplain (2013) took a spatial approach to investigating the roles of membrane-bound MMPs like MT1-MMP and of soluble MMPs like MMP-2.

*Local* models relying on *discrete* and *hybrid* approaches have also been proposed. For instance, individual-based models (IBMs) in (Anderson and Chaplain, 1998; Anderson et al, 2000; Zhang et al, 2009), cellular automata models in (Kansal et al, 2000; Deutsch and Dormann, 2005; Hatzikirou and Deutsch, 2008; Enderling et al, 2009), cellular Potts model approaches in (Turner and Sherratt, 2002; Popławski et al, 2009; Scianna et al, 2013), and hybrid-discrete continuum models in (Anderson, 2005; Rejniak and Anderson, 2011).

Further, *non-local* PDE models in the form of integro-differential equations, which incorporate cell-cell adhesion using integral terms, have been developed. A continuum model of adhesion forces and their influence on cell movement was proposed by Armstrong et al (2006). They accounted for adhesion by an integral term, which modelled non-local interactions in the PDE model. Their model was the first of its kind to include cell-cell adhesion in a continuum model of interacting cell populations. Two years later, Gerisch and Chaplain (2008) based their first non-local cancer invasion PDE model on (Armstrong et al, 2006). They represented one or more cancer cell populations by a PDE each and additionally considered a PDE to represent the ECM, which was modelled to be fixed in space. In this way, cancer cell-cell and cancer cell-matrix adhesion were modelled. Sherratt et al (2009) proposed a similar non-local PDE model of cancer invasion based on the same work by Armstrong et al (2006). Chaplain et al (2011) studied the nature of the proliferative properties of non-local PDE models analytically by proving some results on the basis of the paper by Gerisch and Chaplain (2008). Furthermore, they provided computational simulations illustrating the relative effects of cell-cell and cell-matrix adhesion on cancer invasion. Domschke et al (2014) further developed the model by Gerisch and Chaplain (2008) to study the influence of cell-cell and cell-matrix adhesion on tumour growth and development in more depth. In particular, they introduced a subpopulation of cancer cells of a second type that was assumed to have grown to become more aggressive over time through mutations. A non-local multiscale PDE model for glioma invasion, which considered the clinically observed migration pathways of invading cancer cells along neural fibre tracts, was presented by Painter and Hillen (2013). This model made it possible to study the impact of fibre alignment on the cancer cells’ invasion pathways and to connect diffusion tensor imaging data to the model’s parameters.

Metastasis involves a variety of sub-processes at multiple temporal and spatial scales, as discussed in Section 2. Hence, different models are appropriate to shed light on the various sub-processes. These models can broadly be categorised into those describing how successful metastatic phenotypes evolve by epigenetic and genetic mutations — such as Moran process models by Michor et al (2006) or by Michor and Iwasa (2006), or the time-branching process model by Dingli et al (2007) — and those modelling the growth dynamics of metastases. In what follows, we will give an overview of the latter modelling approaches. Saidel et al (1976) proposed a compartmentalised translational ODE model of metastasis distribution over time. Consecutively, a Markov-Chain model by Liotta et al (1976, 1977) considered a subset of the compartments in (Saidel et al, 1976) to predict metastatic formation. Iwata et al (2000) introduced a hyperbolic PDE model for the colony size distribution of multiple metastatic tumours that form from an untreated tumour, which many subsequent papers used as a basis for their work. For instance, the model by Iwata et al (2000) was further analysed and solved numerically by Barbolosi et al (2009) and Devys et al (2009). It was then used in (Benzekry, 2011) to model metastasis density, while tumour growth and angiogenesis were accounted for by an ODE model by Hahnfeldt et al (1999). The paper by Iwata et al (2000) additionally formed the basis for a mathematical model by Benzekry et al (2016), which connected presurgical primary tumour volume and post-surgical metastatic burden. Finally, an *in vivo* human xenograft model by Hartung et al (2014) was also based on (Iwata et al, 2000). This described primary tumour growth by a set of phenomenological models, and metastatic growth by a transport equation that was endowed with a boundary condition for metastatic emission. Other models of metastatic dynamics include: fully stochastic mechanistic models (Xu and Prorok, 1998; Bartoszyński et al, 2001; Hanin et al, 2006) that used similar growth laws to those proposed by Iwata et al (2000) to predict the probability that a certain given distribution of metastatic colony size occurs at a given time; another stochastic model by Haeno and Michor (2010); and time-branching models by Iwasa et al (2006) and Haeno et al (2007). The latter three are all of similar type to the aforementioned, phenotype-focussed models by Michor et al (2006), Michor and Iwasa (2006) and Dingli et al (2007) but examine an exponentially *expanding* rather than a constant cancer cell population. Out of these three, the models by Iwasa et al (2006) and Haeno et al (2007) studied the dynamics of one or two mutations in metastasis-suppressor genes to investigate the probability that a tumour becomes resistant to therapy. Haeno and Michor (2010) aimed to provide a theoretical example of how to use a mathematical model to examine the effects of the choice of treatment (chemotherapy and/or tumour resection) and of its timing. Scott et al (2013) proposed a model of self-seeding to study the relative likelihood of primary and secondary seeding by assuming that a primary tumour consists of a set of independent loci, on which tumours undergo saturating growth according to a logistic law. From these loci, cancer cells are shed and potentially return to their original loci or form new loci. Newton et al (2013) presented a stochastic Markov Chain/Monte Carlo model to study multidirectional metastatic spread of lung cancer while distinguishing between spreader and sponge metastatic sites, which was extended to be used in breast cancer in (Newton et al, 2015). Cisneros and Newman (2014) proposed another stochastic model that used a birth-death process to investigate whether metastasis occurs from many poorly adapted cancer cells or from a few well-adapted cancer cells. Finally, Margarit and Romanelli (2016) developed a patient-statistics-based absorbing Markov Chain model to analyse the metastatic routes between principal organs.

While all processes in the invasion-metastatic cascade are inherently spatial, we conclude that, to our knowledge, no spatially explicit model to describe metastasis and metastatic spread exists, not to mention a model that combines all of the steps of the invasion-metastasis cascade — i.e. cancer cell invasion, intravasation, vascular travel, extravasation and regrowth at new sites in the body — in a spatial manner. With the aim of closing this gap in the existing literature, we propose a novel spatially explicit hybrid modelling framework that describes the invasive growth dynamics both of the primary tumour and at potential secondary metastatic sites as well as the transport from primary to secondary sites. In what follows, we introduce the ideas and assumptions that our modelling framework builds on. The corresponding modelling algorithm is described in Figure 5 in form of a flowchart.

**Fig. 5:**
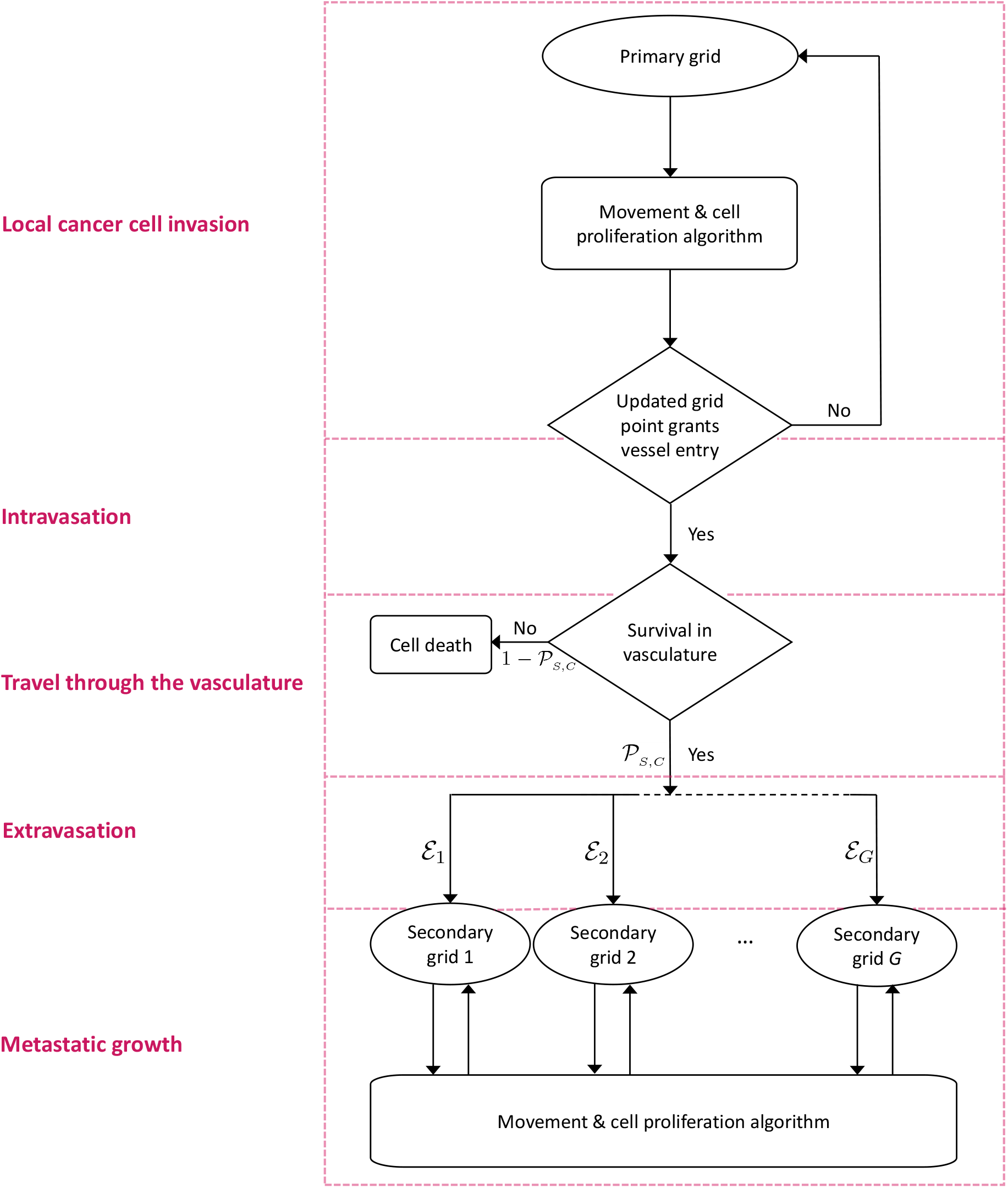
Flowchart of the metastatic algorithm in our hybrid model. At each time step, each cancer cell on the primary grid moves and proliferates according the ‘Movement & cell proliferation algorithm’ explained in detail in the text. A cancer cell remains on the primary grid during the respective time step, unless it is placed on a grid point of the primary grid that represents a blood vessel. In the latter case, single CTCs and CTC clusters may enter the vasculature. They spend some number of time steps in the circulation and survive with a probability of 𝒫_S_ in the case of single CTCs and with a probability of 𝒫_C_ in the case of CTC clusters. Cancer cells that do not survive are removed from the simulation. Surviving CTCs and CTC clusters are placed onto one of G secondary grids with the respective probability Ɛ_1_, Ɛ_2_, …, Ɛ_G_. Cancer cells on the secondary grids perform the same ‘Movement & cell proliferation algorithm’ as cancer cells on the primary grid. For better orientation, the red boxes with their labels on the left correspond to the sections indicated in bold in Section 3 of the text.

In order to account for cancer cell metastasis in a spatially explicit manner, we consider *G* + 1 spatial domains. These consist of the spatial domain representing the primary tumour site, *Ω*_P_, as well as the *G ∈* ℕ spatial domains representing the sites of potential secondary metastatic spread, *Ω^a^_s_*, where *a* = 1, 2, …, *G*. In these spatial domains, we capture the spatiotemporal evolution of *individual* epithelial-like and mesenchymal-like cancer cells, while representing the MMP-2 concentration and the ECM density at position (*x, y*) at time *t* by the continuous functions *m*(*t, x, y*) and *w*(*t, x, y*), respectively. We model the local cancer cell invasion by expanding the modelling approach of Anderson and Chaplain (1998); Anderson et al (2000) to our specific biological problem. We include a second cancer cell phenotype and also additionally consider MT1-MMP, which is taken to be bound to the membranes of the mesenchymal-like cancer cells and thus follows their discrete spatiotemporal dynamics. We designate locations in the primary spatial domain to function as entry points into the vasculature and, similarly, impose a spatial map of exit locations from the vasculature onto the secondary metastatic domains. This allows cancer cells to travel from the primary tumour site to secondary sites via blood vessels.

We next consider one key step of the invasion-metastasis cascade after the other. To make the key steps more recognisable, we begin each paragraph by printing the description of the step in bold. Further, the same step descriptions can be found on the left of the flowchart in Figure 5. This highlights which parts of our algorithm correspond to which sections in the text.

### Local cancer cell invasion

The movement of the individual epithelial-like and mesenchymal-like cancer cells in the spatial domains of our model is *derived* from the coupled PDEs (1) and (2) below. These equations describe the continuous spatio-temporal evolution of epithelial-like and mesenchymal-like cancer cell densities *c*_E_ (*t, x, y*) and *c*_M_ (*t, x, y*), respectively. Both cancer cell types are assumed to move via a combination of diffusive movement and haptotactic movement up the gradient of the ECM density *w*(*t, x, y*). Hence, the evolution of the density of epithelial-like cancer cells *c*_E_ (*t, x, y*) is governed by the following diffusion-haptotaxis equation:

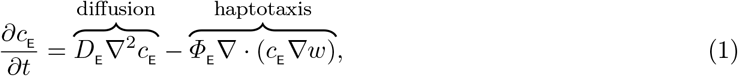

along with zero-flux boundary conditions. Here, *D*_E_ > 0 is the constant cancer cell diffusion coefficient for epithelial-like cancer cells and *Φ*_E_ > 0 is their constant haptotactic sensitivity coefficient. Similarly, the mesenchymal-like cancer cell density *c*_M_ (*t, x, y*) evolves according to:

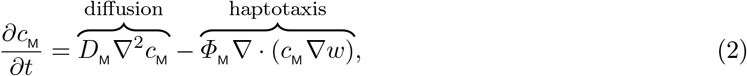

along with zero-flux boundary conditions. Here, *D*_M_ > 0 is the constant cancer cell diffusion coefficient for mesenchymal-like cancer cells and *Φ*_M_ > 0 is their constant haptotactic sensitivity coefficient.

By discretising the spatial domains of our model using a uniform mesh, as described in Appendix B, we derive the movement probabilities of the *individual* epithelial-like and mesenchymal-like cancer cells to be those in equations (B.3). Modelling the cancer cells individually allows us to track the evolution of single mesenchymal-like and epithelial-like cancer cells with different phenotypes and their evolution.

The algorithm we described so far accounts for the movement of our cancer cells only. We thus need to additionally account for the *proliferation* of cancer cells in our model. The two cancer cells types included in our model proliferate at different frequencies. The more proliferative epithelial-like cancer cells perform mitosis after *T*_E_ *∈* ℕ time steps, the less proliferative mesenchymal-like cell types after *T*_M_ *∈* ℕ time steps (with *T*_M_ > *T*_E_). When proliferating, the cancer cells pass on their respective phenotype as well as their location so that a proliferating cancer cell is replace by two daughter cells after the proliferative step has been performed. However, to account for competition for space and resources, the cancer cells on the respective grid point do not proliferate if there are *𝒬 ∈* N cancer cells on a grid point at the time of proliferation.

With reference to the flowchart shown in Figure 5, the part of our approach described so far is summarised in the *Movement & cell proliferation algorithm*, which, for the primary site, is depicted in the upper region of the flowchart.

The mesenchymal-like cancer cells in our model have the ability to express diffusive MMP-2. The MMP-2 concentration *m*(*t, x, y*) hence develops according to the equation:

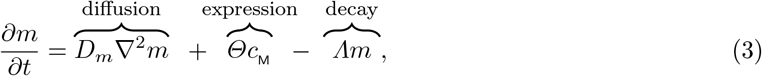

along with zero-flux boundary conditions. Here, *D_m_ >* 0 is the constant MMP-2 diffusion coefficient, *Θ >* 0 is the constant rate of MMP-2 concentration provided by mesenchymal-like cancer cells, and *Λ >* 0 is the constant rate at which MMP-2 decays. Note that the mesenchymal-like cancer cells also express MT1-MMP. However, MT1-MMP acts locally only where it is bound to the cancer cell membrane and its spatiotemporal evolution is hence congruent to that of the mesenchymal-like cancer cells. Therefore, we do not include a separate equation.

The diffusive MMP-2 degrades the ECM with a degradation rate of *Γ*_2_ > 0. The MT1-MMP expressed on the membrane of the mesenchymal-like cancer cells also degrades the ECM, which is expressed through the degradation rate *Γ*_1_ > 0. Hence, given that we are disregarding ECM-remodelling for simplicity, the evolution of the ECM density *m*(*t, x, y*) is governed by the following PDE:

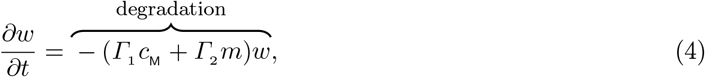

along with zero-flux boundary conditions.

Since the continuous evolution of the MMP-2 concentration and the ECM density is governed by equations 3 and 4, while the spatiotemporal evolution of the cancer cells (and, intrinsically, of the membrane-bound MT1-MMP) is captured by an individual-based model, we model cancer cell invasion in a hybrid-discrete continuum approach. Because the movement probabilities are derived from equations 1 and 2, which are obtained using equations 3 and 4, the hybrid approach is of the kind pioneered by Anderson and Chaplain (1998) to model tumour-angiogenesis, that was subsequently used to model tissue invasion by cancer cells (Anderson et al, 2000; Anderson, 2005) and spatial evolutionary games (Burgess et al, 2016, 2017).

### Intravasation

With the model setup we have described so far, the cancer cells can invade the tissue locally in the primary spatial domain but cannot reach the spatially separated secondary domains. In order to allow for metastatic spread, we account for the connection of the primary spatial domain to the secondary spatial domains by incorporating blood vessels in our modelling framework. Examples of primary and secondary domains are presented in Figure 6. To represent the entry points into the blood vessels, a number of *U*_P_ *∈* ℕ_0_ normal blood vessels as well as *V*_P_ *∈* ℕ_0_ ruptured blood vessels are distributed on the primary grid. The normal blood vessels take the size of one grid point, while ruptured vessels consist of a group of *A^b^ ∈* ℕ, where *b* = 1, 2, …, *V_P_*, adjacent grid points and can thus have different shapes. Each secondary grid *Ω_s_^a^* also has, respectively, *U_s_^a^ ∈* ℕ normal blood vessels, where *a* = 1, 2, …, *G* as before, that take the form of a single grid point each. On the primary grid, the grid points where the vessels are located allow the cancer cells to intravasate, while the respective grid points on the secondary grid allow for extravasation.

**Fig. 6:**
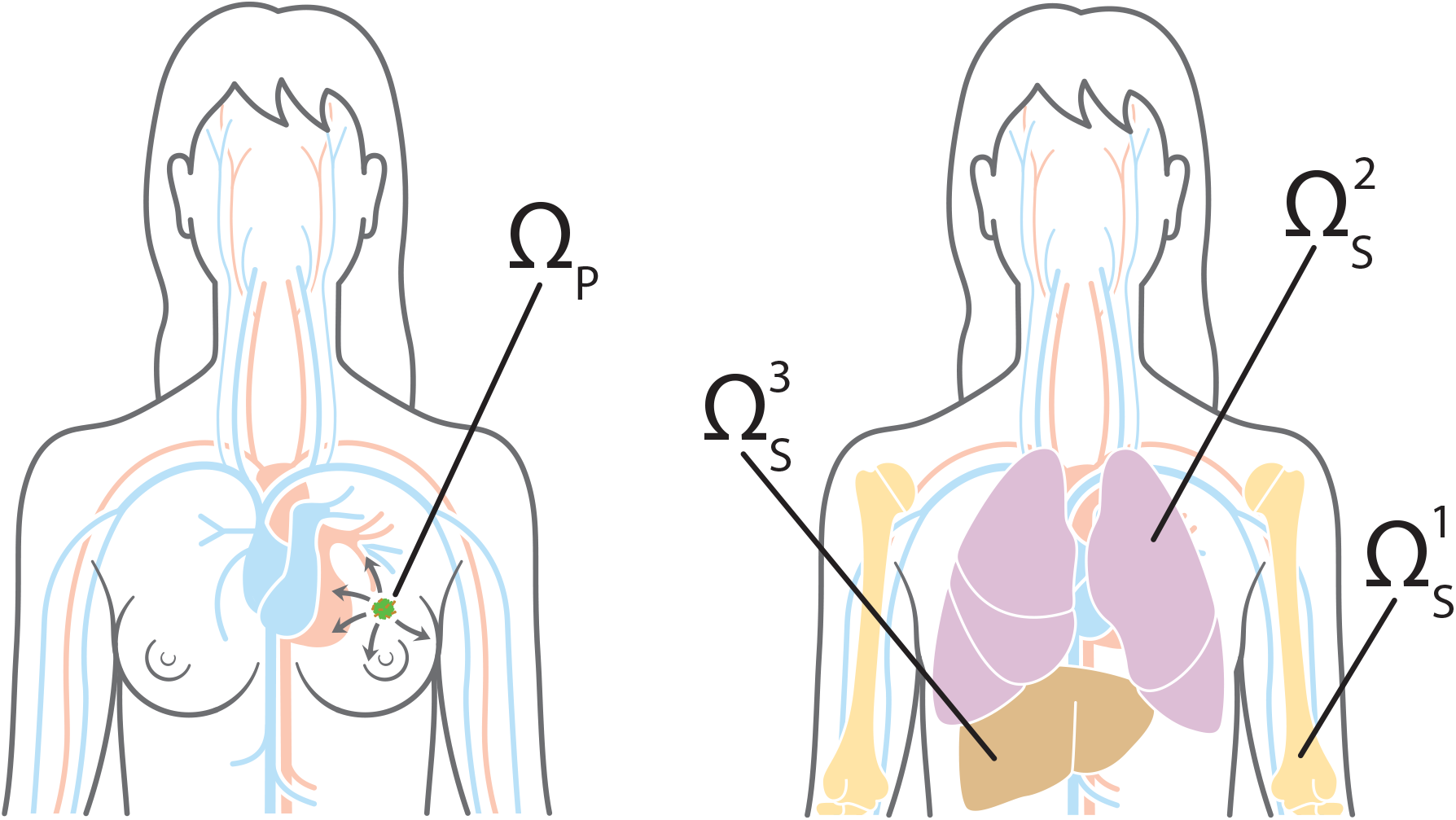
Primary and metastatic sites. To give an example of how our general model can be applied to a specific clinical setting, we chose our primary site Ω_P_, which is shown on the left, to represent the breast, and potential secondary metastatic sites 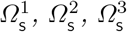, which are shown on the right, to represent the bones, the lungs and the liver, respectively. Cancer cells can reach the secondary sites by travelling through the blood system.

If, by the movement algorithm described in Appendix B, a cancer cell on the primary grid is placed on a grid point that represents a blood vessel, it *may* leave the grid and enter the vasculature. Whether or not a cancer cell can successfully intravasate depends both on its own phenotype and on the type of vessel it is placed on.

Whenever a mesenchymal-like cancer cell is moved to a grid point (*x_i_, y_j_*) *∈ Ω*_P_^1^, on which a *normal* single blood vessel is located, it will successfully enter the vasculature. Further, to represent collective invasion in the form of co-presence of mesenchymal-like and epithelial-like cancer cells, cancer cells of any type on the four neighbouring primary grid points (*x_i_*_+1_, *y_j_*), (*x_i−_*_1_, *y_j_*), (*x_i_, y_j_*_+1_) and (*x_i_, y_j−_*_1_) are forced into the vasculature together with the mesenchymal-like cancer cell on (*x_i_, y_j_*). Hence, a mesenchymal-like cancer cell moving to a grid point on which a normal blood vessel is located results in either a single mesenchymal-like cancer cell or a cluster consisting of up to 5*𝒬* cancer cells of any phenotype intravasating. However, if an epithelial-like cancer cell is moved to a grid point (*x_i_, y_j_*) *∈ Ω*_P_ where a *normal* single vessel is located without a mesenchymal-like cell being present there, the epithelial-like cancer cell will not intravasate and the grid point (*x_i_, y_j_*) will be treated like any other grid point. This is to model the fact that epithelial-like cancer cells have been shown to be unable to actively intravasate on their own.

Further, a cancer cell on the primary grid can move to one of the grid points where a *ruptured* vessel is located. Contrary to the above-described scenario of entering a normal vessel, a cancer cell of any type, which is placed on a grid point representing part of a ruptured vessel, can enter the circulation. The respective cancer cell takes with it any other cancer cells residing both on the grid points representing the ruptured blood vessel and on the regular grid points bordering the ruptured vessel. Biologically, the fact that cancer cells of any phenotype can intravasate mirrors that these blood vessels are already ruptured due to trauma or pressure applied by the expanding tumour, making the requirement of MDE-mediated degradation of the vessel wall redundant. The fact that other cancer cells on bordering grid points will enter the circulation together with cancer cells placed on grid points representing blood vessels captures some degree of the cell-cell adhesion found in collectively invading cancer cell clusters.

### Travel through the vasculature

If a cancer cell of either phenotype or a cluster of cancer cells successfully enter the vasculature either through a ruptured or a normal vessel, it will be removed from the primary grid and moved to the vasculature. Cancer cells and cancer cell clusters remain in the vasculature for some number of time steps *T_V_ ∈* ℕ, which biologically represents the average time the cancer cells spend in the blood system. Any cancer cells that enter a particular vessel at the same time are treated as one cluster and hence as a single entity once they are located in the vasculature. However, each cancer cell that is part of a cancer cell cluster disaggregates from its cluster with some probability *𝒫_d_* after 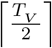, time steps. At the end of *T_V_* time steps, the single cancer cells and the remaining cancer cell clusters are removed from the simulation unless they are randomly determined to survive. The survival probability is *𝒫_S_* for single cancer cells and *𝒫_C_* for cancer cell clusters.

### Extravasation

Any surviving cancer cells and cancer cell clusters are placed on one of the *G* secondary grids *Ω^a^ _S_* with probability *Ɛ*_1_, *Ɛ*_2_, …, *Ɛ_G_*, where 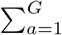 *Ɛ_a_* = 1. Also, on each specific secondary grid, the cancer cells extravasate through one of the randomly chosen *U_S_^a^* grid points that represent a blood vessel with equal probability. If the respective grid point cannot accommodate all of the entering cancer cells without violating the carrying capacity *Q*, the remaining cancer cells are randomly distributed onto the four non-diagonally neighbouring grid points until these are filled to carrying capacity *𝒬*. If there are further cancer cells to be placed onto the respective grid point at this instance, such cancer cells are killed to capture the influence of competition for space in combination with vascular flow dynamics.

### Metastatic growth

If and when cancer cells reach a secondary grid, they behave (i.e. replicate, move, produce MDEs etc.) there according to the same rules as on the primary grid, as indicated on the bottom of the flowchart in Figure 5.

## 4 Setup of computational simulations and model calibration

To perform our numerical simulations, we non-dimensionalised the system of equations (1)–(4) as described in Appendix A. Like Anderson et al (2000), we choose to rescale distance with an appropriate length scale *L* = 0.2 cm (since 0.1–1 cm is estimated to be the maximum invasion distance of cancer cells at an early stage of cancer invasion) and time with an appropriate scaling parameter 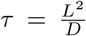. Here *D* = 10^−6^ cm^2^s^−1^ is a reference chemical diffusion coefficient suggested by Bray (1992), such that *τ* = 4 × 10^4^ s, which corresponds to approximately 11 h.

We considered spatial domains of size [0, 1] × [0, 1], which corresponds to physical domains of size [0, 0.2]cm × [0, 0.2]cm. In particular, we let the spatial domain *Ω*_P_ represent the primary site and the spatial domains 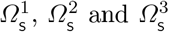 describe three metastatic sites. These spatial domains could represent *any* primary and secondary carcinoma sites. However, to give an example of a particular application, we considered a study of 4181 breast cancer patients at Memorial Sloan Kettering Cancer Center. Data and graphs from this study can be found at http://kuhn.usc.edu/breast_cancer (Kuhn Laboratory, 2017). We accordingly chose *Ω*_P_ to represent the primary site of the breast, and 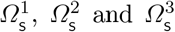 to correspond to the common metastatic sites of bones, lungs and liver, respectively. Disregarding potential spread to any other metastatic sites, the data from (Kuhn Laboratory, 2017) provided us with an extravasation probability of *Ɛ*_1_ *≈* 0.5461 to the bones, of *Ɛ*_2_ *≈* 0.2553 to the lungs, and of *Ɛ*_3_ *≈* 0.1986 to the liver.

We discretised the four spatial domains to contain 201 × 201 grid points each. This corresponds to a non-dimensionalised space step of *Δx* = *Δy* = 5 × 10^−3^, which results in a dimensional space step of 1 × 10^−3^ cm, and thus roughly corresponds to the diameter of a breast cancer cell (Vajtai, 2013). We then chose a time step of *Δt* = 1 × 10^−3^, corresponding to 40 s, to comply with the Courant-Friedrichs-Lewy condition (Anderson et al, 2000). We ran our simulation for 48000*Δt* time steps, which corresponds to *∼*22 days.

On each secondary grid, we chose 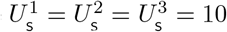 distinct grid points, on which blood vessels are located. For each grid, these blood vessels were placed randomly but at least two grid step widths away from the respective grid’s boundary. The same applies to the primary grid *Ω*_P_ but with the additional condition that the *U*_P_ = 8 single grid points, where normal blood vessels are located, and the *V*_P_ = 2 sets of five grid points, where ruptured blood vessels are placed, are located outside a quasi-circular region containing the 200 centre-most grid points. While these 10 randomly placed vessels are modelled to exist from the beginning, they represent those vessels that grow as a result of tumour-induced angiogenesis in the vascular tumour growth phase.

To represent a two-dimensional cross-section of a small avascular primary tumour, we placed a nodule that consisted of 388 randomly distributed cancer cells in the quasi-circular region of the 97 centre-most grid points of the primary grid, with no more than *𝒬* = 4 cancer cells on any grid point to account for competition for space. This carrying capacity of *𝒬* = 4 was applied throughout the simulation. A randomly chosen 40% of these cancer cells were of epithelial-like phenotype, and the remaining 60% of mesenchymal-like phenotype. The described initial condition ensures that the cancer cells are placed away from any pre-existing vessels to match the biology of an avascular tumour. Figure 7 gives an example of a typical initial cancer cell placement and vessel distribution on the primary grid.

**Fig. 7:**
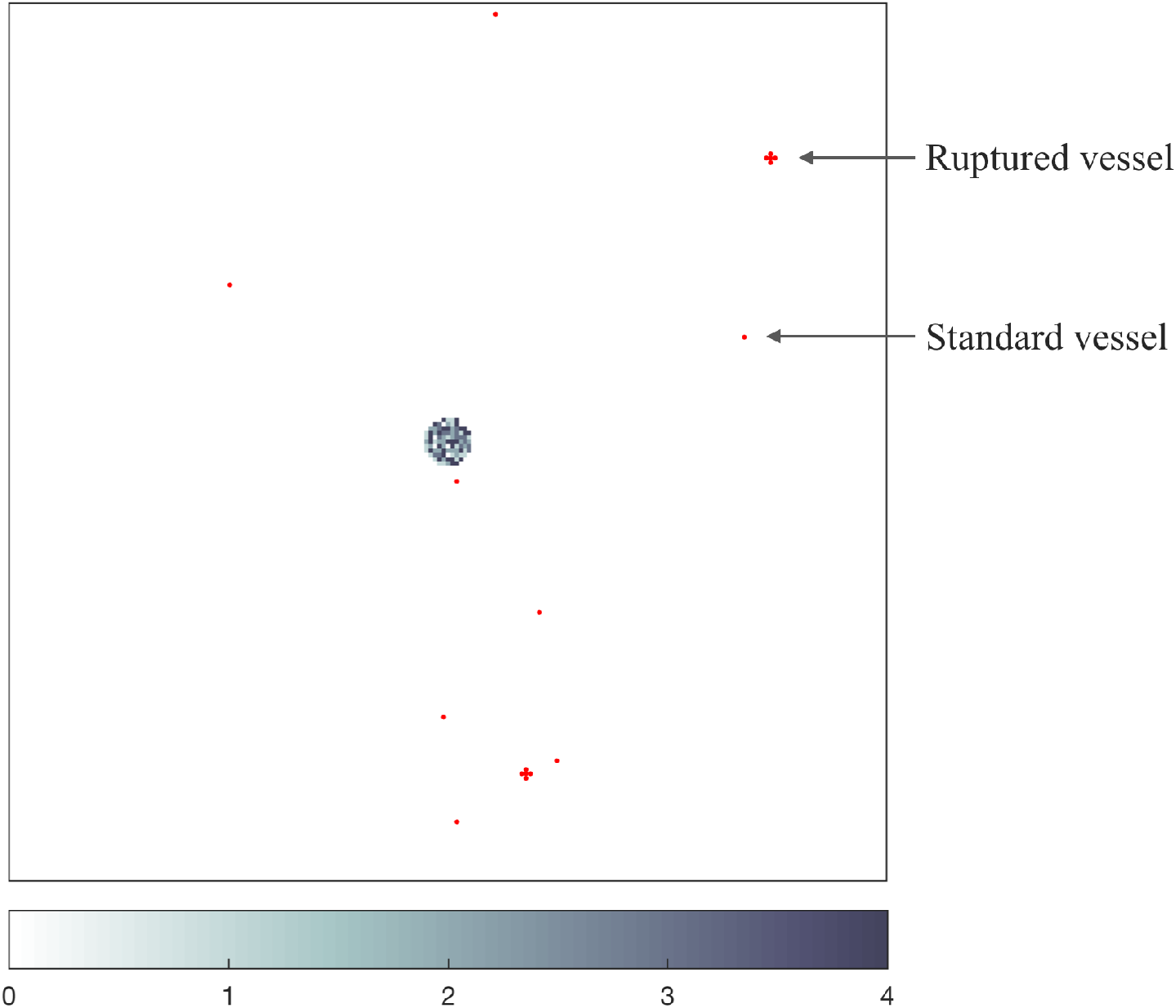
Vessel distribution and initial condition of cancer cells. The plot shows ten randomly distributed blood vessels on the primary grid, two of which are so-called ruptured vessels that consist of five rather than one grid point. In the centre of the grid, the initial cancer cell distribution is shown. There are between 1 (light grey) and 4 (black) cancer cells on a grid point. As the initial distribution of cancer cells represents a 2D cut through an avascular tumour, the blood vessels are placed at some distance away from the initial nodule of cancer cells.

In accordance with the ranges provided in Table 1, we chose the mesenchymal-like cancer cell diffusion coefficient to be *D_M_* = 1 × 10^−4^, the epithelial-like cancer cell diffusion coefficient to be *D_E_* = 5 × 10^−5^, and the mesenchymal and epithelial haptotactic sensitivity coefficients to be *Φ_M_* = *Φ_M_* = 5 × 10^−4^.

**Table 1:** Baseline parameter settings for our simulations. In the first column, non-dimensional parameters are indicated by upper-case notation. Corresponding dimensional parameters are stated in brackets using lower-case notation. In the fourth column, we reference other mathematical modelling papers in brackets and biological papers without brackets.

|  | Description | Non-dimen-<br>sional value | Biological reference<br>(Modelling reference) | Original value |
| --- | --- | --- | --- | --- |
| $\Delta t$ | Time step | $1 \times 10^{-3}$ | | 40 s |
| $\Delta x,$<br>$\Delta y$ | Space step | $5 \times 10^{-3}$ | Breast cell diameter in<br>Vajtai (2013) | $1 \times 10^{-3}$ cm |
| $D_M(d_M)$ | Mesenchymal-like cancer<br>cell diffusion coefficient | $1 \times 10^{-4}$ | Bray (1992)<br>(Anderson et al (2000))<br>(Deakin and Chaplain (2013)) | $1 \times 10^{-10}$ cm<sup>2</sup>s<sup>-1</sup> |
| $D_E(d_E)$ | Epithelial-like cancer<br>cell diffusion coefficient | $5 \times 10^{-5}$ | Bray (1992)<br>(Anderson et al (2000))<br>(Deakin and Chaplain (2013) ) | $5 \times 10^{-11}$ cm<sup>2</sup>s<sup>-1</sup> |
| $\Phi_M(\phi_M)$ | Mesenchymal haptotactic<br>sensitivity coefficient | $5 \times 10^{-4}$ | Stokes et al (1990)<br>(Anderson et al (2000)) | $2.6 \times 10^3$ cm<sup>2</sup>M<sup>-1</sup>s<sup>-1</sup> |
| $\Phi_E(\phi_E)$ | Epithelial haptotactic<br>sensitivity coefficient | $5 \times 10^{-4}$ | Stokes et al (1990)<br>(Anderson et al (2000)) | $2.6 \times 10^3$ cm<sup>2</sup>M<sup>-1</sup>s<sup>-1</sup> |
| $D_m(d_m)$ | MMP-2 diffusion<br>coefficient | $1 \times 10^{-3}$ | Collier et al (2011) | $1 \times 10^{-9}$ cm<sup>2</sup>s<sup>-1</sup> |
| $\Theta(\theta)$ | MMP-2 production rate | 0.195 | Estimated | $4.875 \times 10^{-6}$ Ms<sup>-1</sup> |
| $\Lambda(\lambda)$ | MMP-2 degradation rate | 0.1 | Estimated in<br>(Deakin and Chaplain, 2013) | $2.5 \times 10^{-6}$ s<sup>-1</sup> |
| $\Gamma_1(\gamma_1)$ | ECM degradation<br>rate by MT1-MMP | 1 | Based on<br>(Deakin and Chaplain, 2013) | $1 \times 10^{-4}$ s<sup>-1</sup> |
| $\Gamma_2(\gamma_2)$ | ECM degradation<br>rate by MMP-2 | 1 | Based on<br>(Anderson et al, 2000) | $1 \times 10^{-4}$ M<sup>-1</sup>s<sup>-1</sup> |
| $T_V$ | Time CTCs spend in the<br>vasculature | 0.18 | Meng et al (2004) | $7.2 \times 10^3$ s |
| $T_M$ | Epithelial doubling time | 3 | Milo et al (2009) | $1.2 \times 10^5$ s |
| $T_E$ | Mesenchymal doubling<br>time | 2 | Milo et al (2009) | $8 \times 10^4$ s |
| $\mathcal{P}_S$ | Single CTC survival<br>probability | $5 \times 10^{-4}$ | Luzzi et al (1998) | $5 \times 10^{-4}$ |
| $\mathcal{P}_C$ | CTC cluster survival<br>probability | $2.5 \times 10^{-2}$ | Luzzi et al (1998)<br>Aceto et al (2014) | $2.5 \times 10^{-2}$ |
| $\mathcal{E}_1$ | Extravasation probability<br>to bones | $\sim 0.5461$ | Kuhn Laboratory (2017) | $\sim 0.5461$ |
| $\mathcal{E}_2$ | Extravasation probability<br>to lungs | $\sim 0.2553$ | Kuhn Laboratory (2017) | $\sim 0.2553$ |
| $\mathcal{E}_3$ | Extravasation probability<br>to liver | $\sim 0.1986$ | Kuhn Laboratory (2017) | $\sim 0.1986$ |

We further assumed that, once in the vasculature, a single CTC had a survival probability of *𝒫_S_* = 5 × 10^−^, which is of the order of the micro- and macrometastatic growth success rates proposed in (Luzzi et al, 1998). We chose the success rate for metastatic growth to be our survival probability because our model in its current state disregards cancer cell death at secondary sites so that any successfully extravasated cancer cell will initiate micrometastatic growth over time. CTC clusters had a survival probability *𝒫_C_* = 50*𝒫_S_* = 2.5 × 10^−^, in accordance with the finding by Aceto et al (2014) that the survival probability of CTC clusters is between 23 and 50 times higher than that of single CTCs. Surviving single CTCs and CTC clusters exited onto the secondary grids after spending *T_V_* = 0.18 in the blood system, which corresponds to 2 h and hence to the breast cancer-specific clinical results in (Meng et al, 2004).

Further, we assumed a uniform initial ECM density of *w*(*t, x, y*) = 1 across all the spatial domains, while the initial MMP-2 concentration was *m*(*t, x, y*) = 0. We chose the other parameters as shown in Table 1 and assumed that epithelial-like cancer cells divide by mitosis every *T*_E_ = 2000 time steps and mesenchymal-like cancer cells every *T*_M_ = 3000 time steps. This corresponds to approximately 22 hours and 33 hours, respectively, which is consistent with the average doubling times found in breast cancer cell lines (Milo et al, 2009).

## 5 Computational simulation results

To verify that our modelling framework is suitable to capture the key steps of the invasion-metastasis cascade, we first ran simulations with the base parameters shown in Table 1. As indicated by the headings throughout this section, we then varied these base parameters across biologically realistic ranges to further confirm that our framework delivers biologically realistic results and to gain insight into the underlying biology. For each of the parameter studies, we took the average results from running the simulation three times — unless stated otherwise — and indicated error bars to display the standard deviation in the corresponding plots. This way, we studied the effect that changing the initial ratio of epithelial-like to mesenchymal-like cancer cells, the number of blood vessels in the primary site, and the survival probability had on the overall cancer dynamics. We investigated the roles of MMP-2 and MT1-MMP as well as their role in comparison to one another. Finally, we changed the parameters to describe haptotaxis-dominated rather than diffusion-dominated cancer cell movement at the end of this section to re-examine the role of membrane-bound versus diffusive MDEs. We compared the outcomes of these simulations to a range of experimental and clinical results.

## Simulations with base parameters

When using the settings outlined in the previous section, we observed in our simulations that both epithelial-like and mesenchymal-like cancer cells invaded the tissue surrounding the primary tumour, which is represented by the primary grid, over a 22 day period. This is shown in the simulation results in the two upper rows of panels in Figure 8, respectively. The epithelial-like cancer cells formed the bulk of the central tumour mass, while the mesenchymal-like cancer cells were predominantly found at the outermost tissue-invading edge. The maximum observed invasion distance of the cancer cells over this time span was approximately 0.13 cm. The pattern of MMP-2 concentration for the same simulation roughly followed the distribution of the mesenchymal-like cancer cells as shown in the third row of panels in Figure 8. The ECM density, which is depicted in the bottom row of Figure 8, also followed the evolution of the MMP-2 concentration but in a more uniform fashion.

**Fig. 8:**
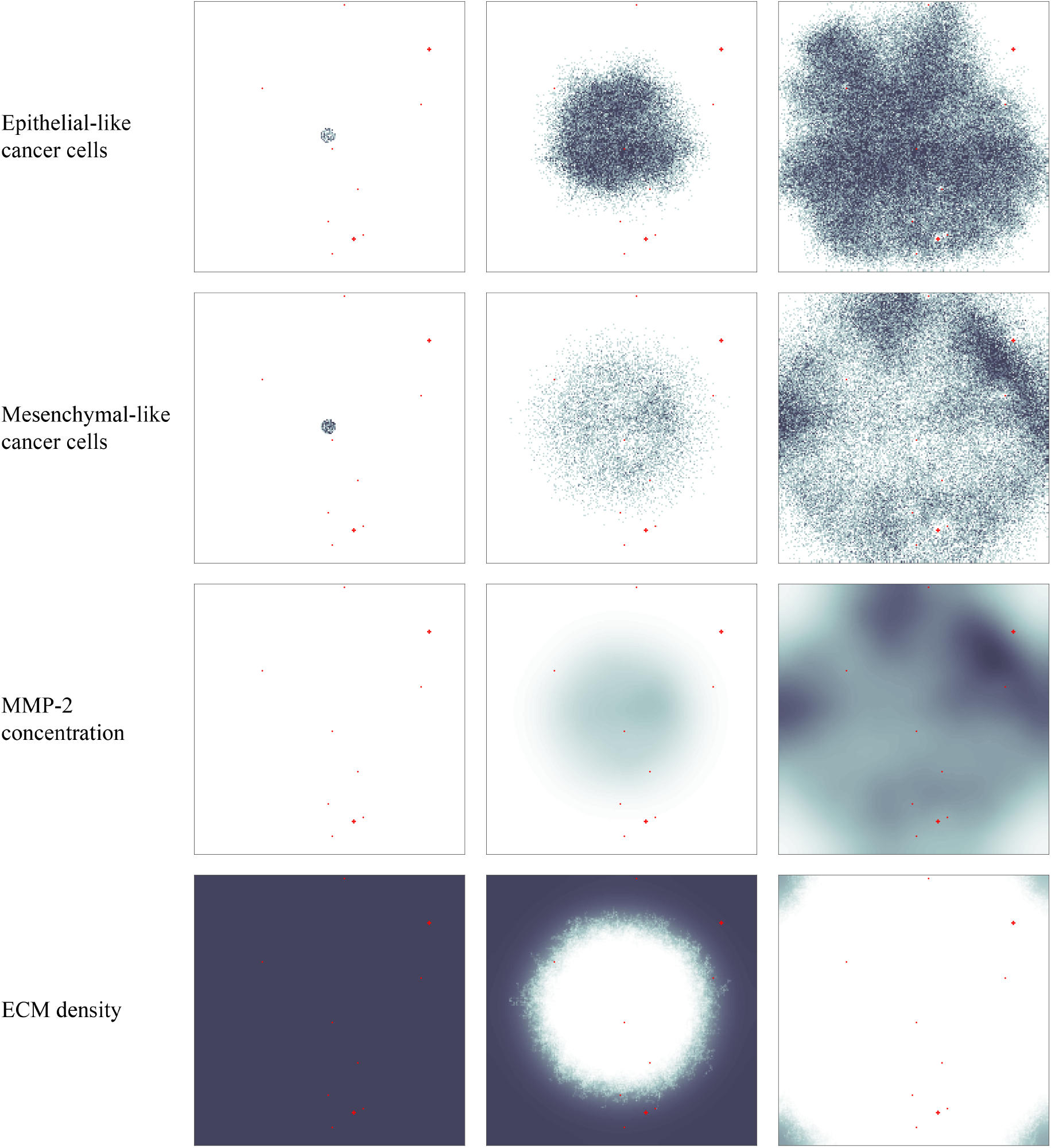
Simulation results on the primary grid. Primary tumour dynamics at times 0, 24000Δt and 48000Δt (left to right), corresponding to 0 days, ∼11 days and ∼22 days. For each time step, the distribution of epithelial-like cancer cells (top row) and mesenchymal-like cancer cells (second row) is shown, with the discrete number of cancer cells per grid point ranging from 1 (light grey) to 4 (black) on each of the panels. The MMP-2 concentration (third row) continuously varies between 0 (white) and 3.0936 (black), and the ECM density (bottom row) between 0 and 1. Red dots represent blood vessels. There are 8 normal blood vessels of the size of one grid point as well as 2 ruptured blood vessels, which extend over 5 grid points each. If cancer cells are moved to these grid points that represent blood vessels, they may enter the vasculature and can potentially extravasate at secondary sites. The dynamics of the cancer cells at the secondary sites are presented in Figures 9–11.

In addition to the cancer cell invasion on the primary grid, we also observed metastatic spread of single cancer cells, as well as homogeneous and heterogeneous cancer cell clusters, to the grids representing the secondary sites of the bones (Figure 9), the lungs (Figure 10) and the liver (Figure 11). The results obtained here showed that the first metastatic spread occurred at the site of the bones. As shown in the panel on the top left of Figure 9, after 11 days we already observed a micrometastatic lesion of epithelial-like cancer cells with an approximate diameter of 0.04 cm on the grid that represented the bones, but in none of the other locations. Yet, after 22 days we discovered metastatic spread at all three of the secondary locations in the body that we considered in our simulations. On the grid representing the bones, we found that the earliest micrometastasis, which consisted of epithelial-like cancer cells only, had rapidly increased in diameter to approximately 0.1 cm (see top right panel of Figure 9). Additionally, a second micrometastasis consisting of both epithelial-like and mesenchymal-like cancer cells had formed at the top right of the same grid. Finally, we observed a set of four epithelial-like and two mesenchymal-like DTCs at the bottom right of the grid corresponding to the bones. The latter two formations were results of heterogeneous cancer cell clusters spreading to the bones. While the secondary site that represented the bones showed by far the largest cancer cell load after 22 days, we detected a further two micrometastases at the secondary site of the lungs, as shown in the panels on the right of Figure 10. While the micrometastasis at the bottom of the grid consisted of epithelial-like cancer cells only, the top micrometastasis contained both cancer cell types as it had grown out of a heterogeneous cancer cell cluster. Finally, the liver showed the least secondary spread in the form of three mesenchymal-like DTCs that arrived as a single cluster, as shown in the panel on the bottom right of Figure 11.

**Fig. 9:**
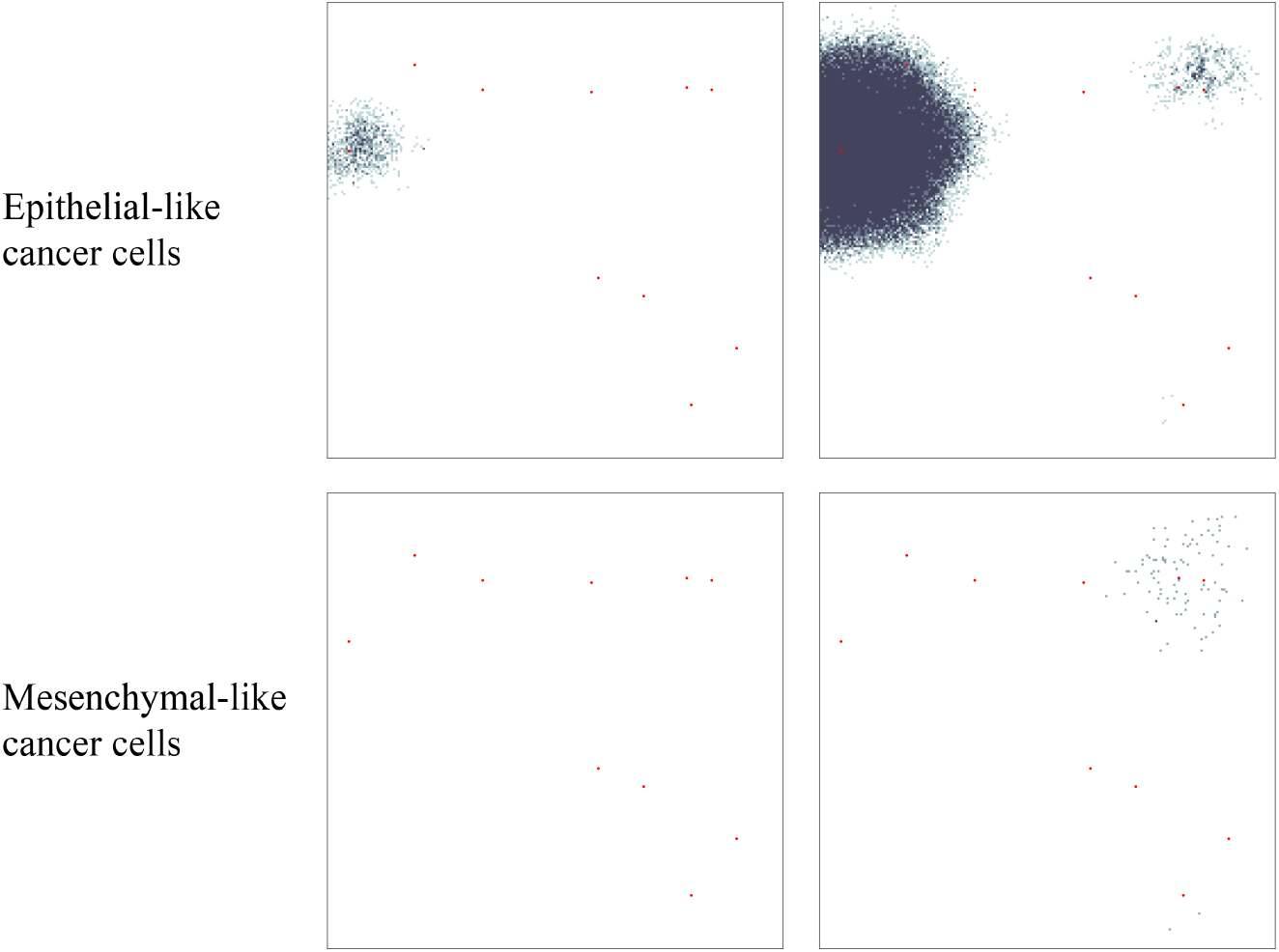
Simulation results on secondary grid representing the bones. Distribution of epithelial-like cancer cells (upper panels) and mesenchymal-like cancer cells (lower panels) at the secondary site representing the bones is shown at times 24000Δt (left) and 48000Δt (right), which corresponds to ∼11 days and ∼22 days. The number of cancer cells per grid point varies between 1 (light grey) and 4 (black) in the upper panels and between 1 (grey) and 2 (black) in the lower panels. The corresponding MMP-2 concentration and ECM density plots are presented in Figure 18 of Appendix C.

**Fig. 10:**
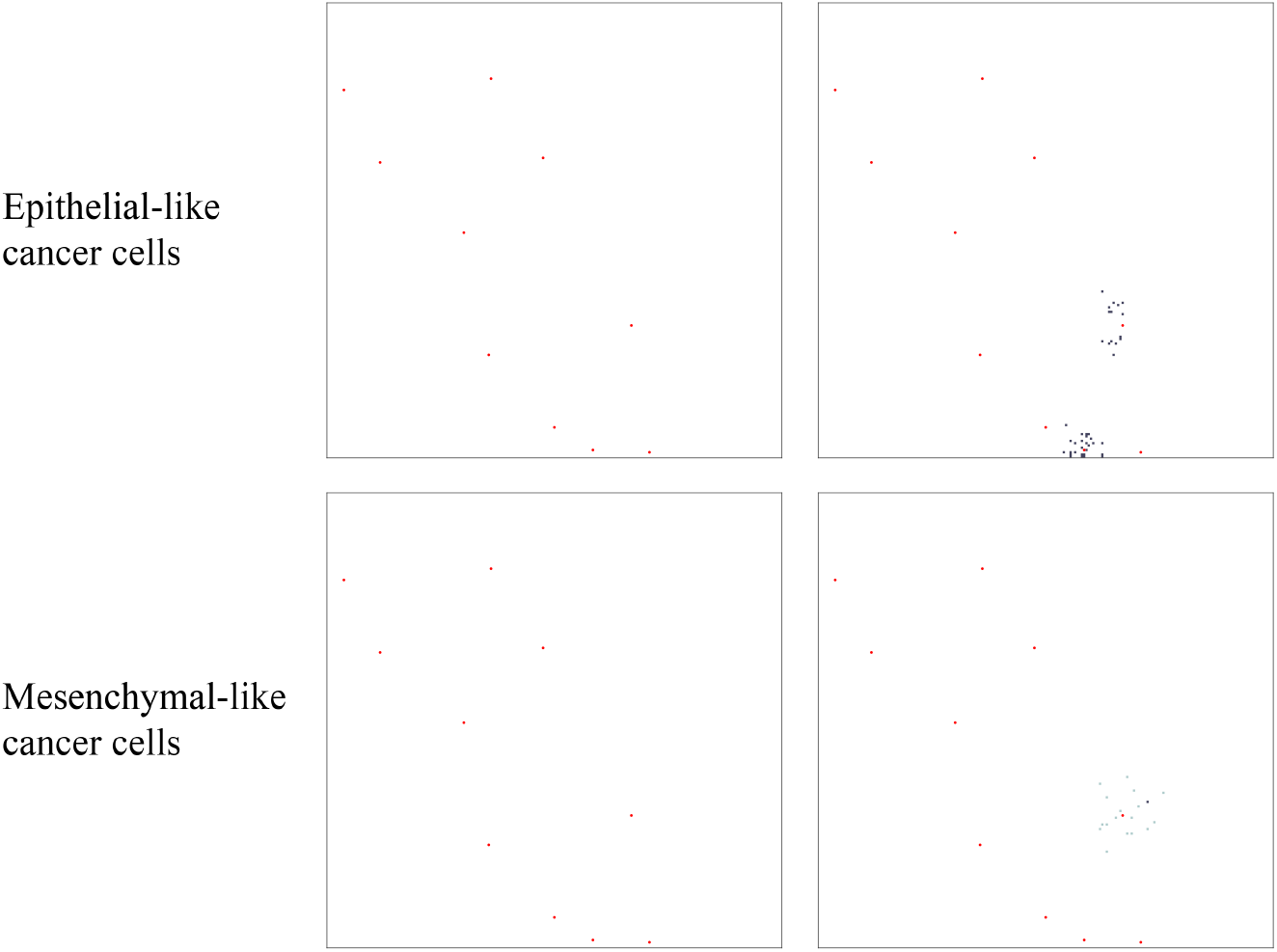
Simulation results on secondary grid representing the lungs. Distribution of epithelial-like cancer cells (upper panels) and mesenchymal-like cancer cells (lower panels) at the secondary site representing the lungs is shown at times 24000Δt (left) and 48000Δt (right), which corresponds to ∼11 days and ∼22 days, respectively. The number of cancer cells per grid point varies between 1 (grey) and 2 (black) in the upper panels and 1 (light grey) and 3 (black) in the lower panels. The corresponding MMP-2 concentration and ECM density plots are presented in Figure 19 of Appendix C.

**Fig. 11:**
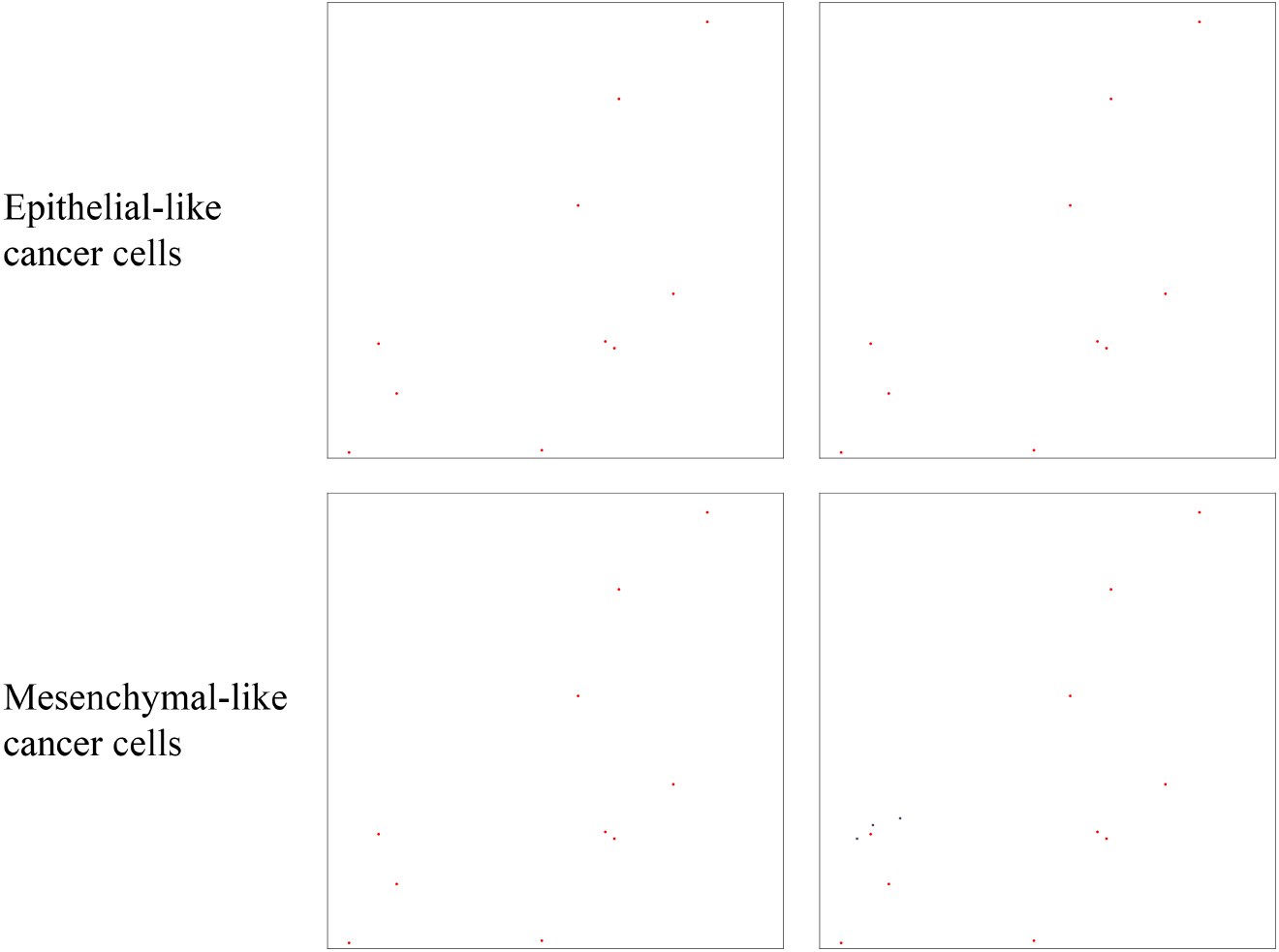
Simulation results on secondary grid representing the liver. Distribution of epithelial-like cancer cells (upper panels) and mesenchymal-like cancer cells (lower panels) at the secondary site representing the liver is shown at times 24000Δt (left) and 48000Δt (right), which corresponds to ∼11 days and ∼22 days, respectively. The panel on the bottom right contains three single mesenchymal-like cancer cells indicated in black, while the other panels do not contain any cells. The corresponding MMP-2 concentration and ECM density plots are presented in Figure 20 of Appendix C.

## Changing the initial ratio of epithelial- to mesenchymal-like cancer cells

By keeping the total initial amount of cancer cells constant at 388 but varying the initial percentage of epithelial-like cancer cells between 0% and 100% in steps of 10%, we found that having solely epithelial-like cells at the start of the simulation had a significant negative impact on tumour growth. We counted an average of 62932 cancer cells at the end of our simulation timespan of 22 days as compared to about 48% more cancer cells (i.e. 93115 cancer cells) in the case of an even initial distribution.

Starting the simulation solely with mesenchymal-like cancer cells had a similar, yet weaker, dampening effect: Compared to simulation with an even initial distribution, it reduced growth to 86425, and thus by about 7.7%. Otherwise, we generally found that a higher percentage of epithelial-like cancer cells at the start coincided with a lower number of mesenchymal-like cancer cells at the end of the simulations. At the same time, the number of epithelial-like cancer cells after 22 days increased. We observed that the maximum number of cancer cells occurred under initial conditions with even parts of mesenchymal-like and epithelial-like cancer cells but that the combined cancer cell count at the end of the simulation was relatively stable if we varied the initial number of epithelial-like cancer cells between 0% and 90% (see Figure 12).

**Fig. 12:**
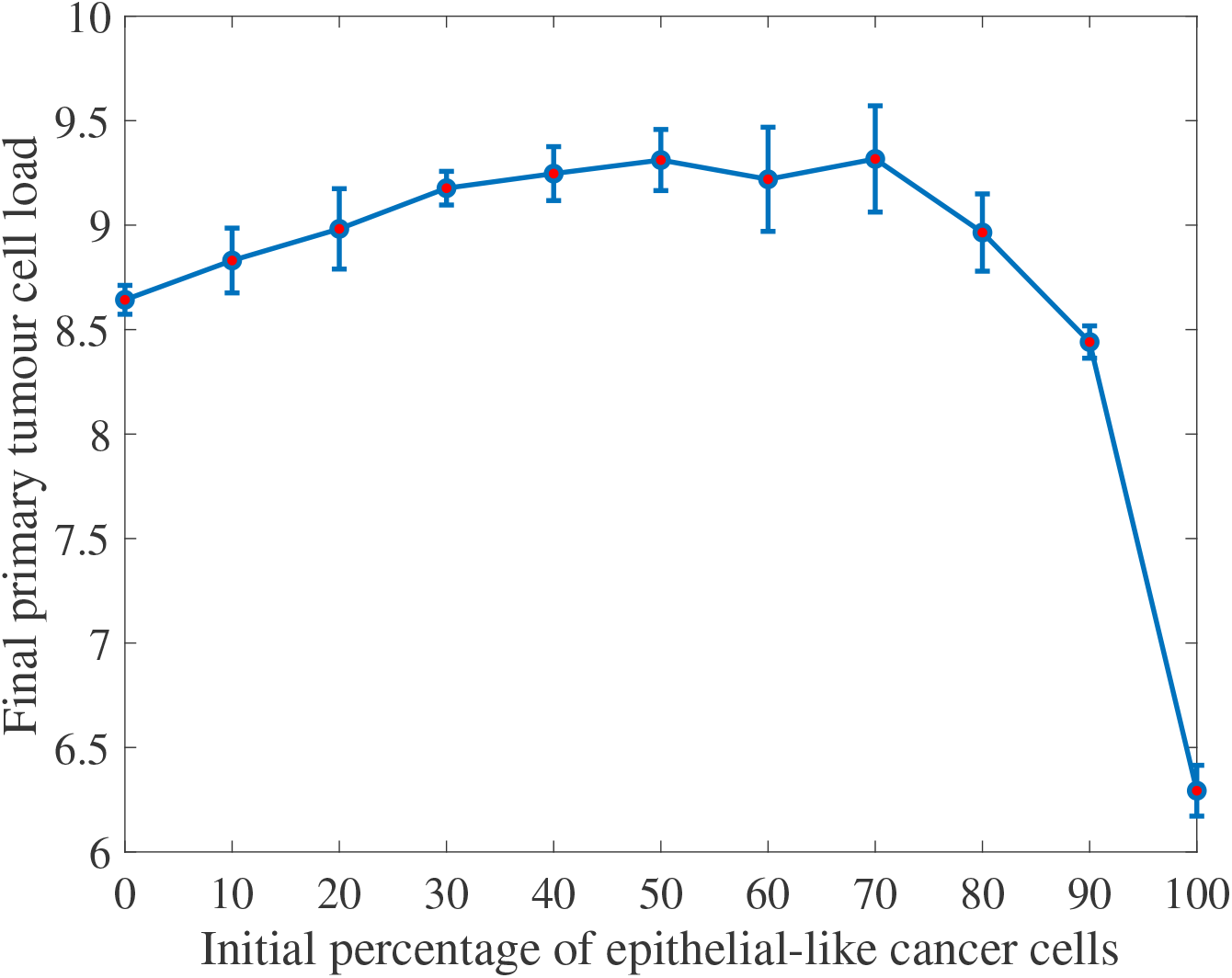
Co-presence of epithelial- and mesenchymal-like cancer cells increases the overall primary tumour cancer cell load. The absence of mesenchymal-like cancer cells hinders cancer cell invasion and tumour growth. The final primary tumour cancer cell load on the vertical axis is given in units of 10^4^ and refers to simulation results after approximately 22 days.

With regards to shedding from the primary tumour, and hence also to chances of successful metastasis, we found that a higher initial percentage of mesenchymal-like cancer cells correlated to a higher number of intravasating single cancer cells and cancer cell clusters, likely as a result of an overall higher number of mesenchymal-like cancer cells (see Figure 13). If we started our simulation with mesenchymal-like cancer cells only, we observed an average total of 634 intravasations by single cancer cells or cell clusters — compared to only 7 over the same time range in the case of the average of simulations that included epithelial-like cancer cells only. When we set the number of ruptured vessels in the primary grid to 0 and considered 10 normal vessels only, we observed no intravasations.

**Fig. 13:**
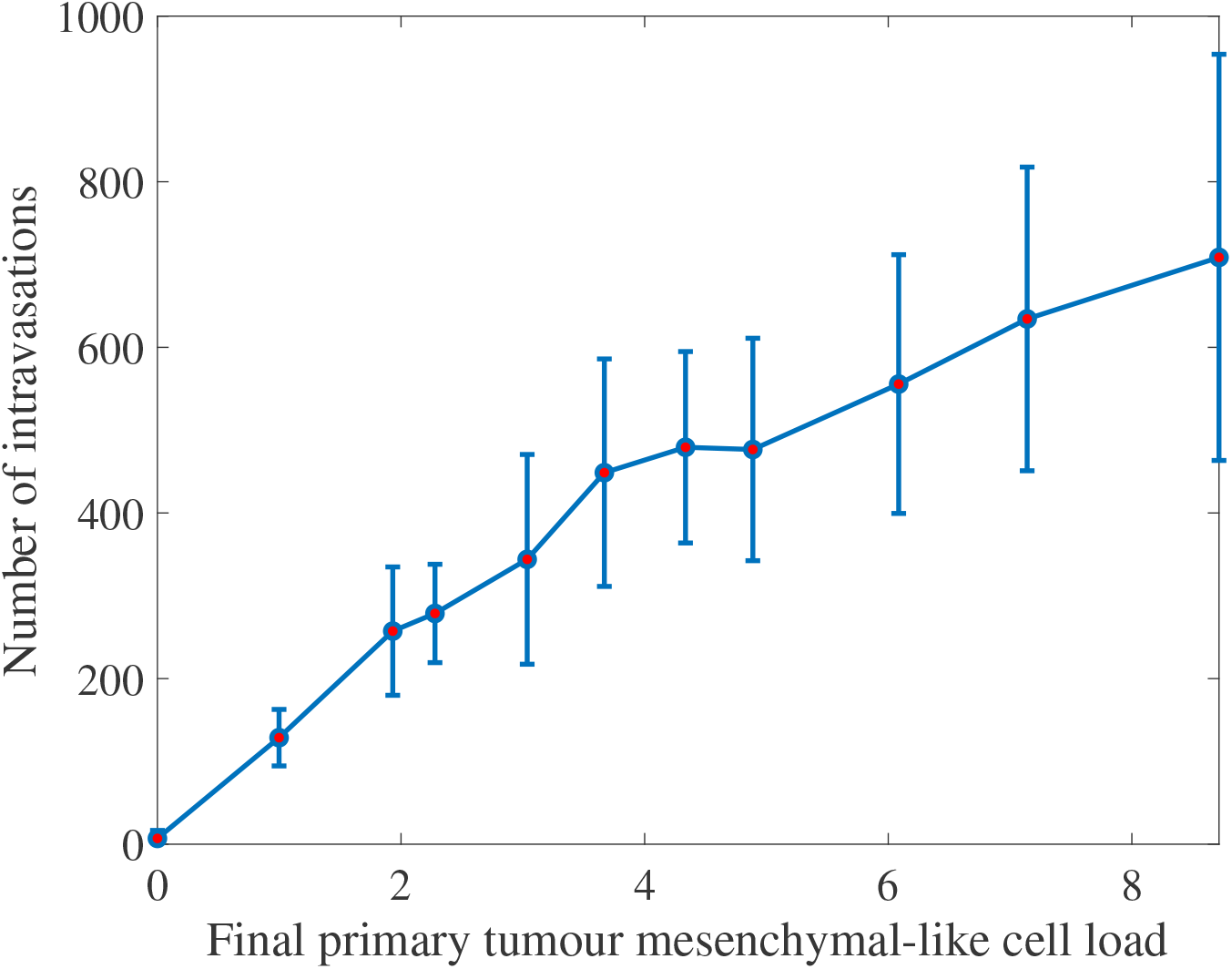
Higher numbers of mesenchymal-like cancer cells at the primary site correspond to an increased intravasation count. The final primary tumour cancer cell load on the horizontal axis is given in units of 10^4^ and refers to simulation results after approximately 22 days.

## Changing the survival probability of cells in the vasculature

As Aceto et al (2014) suggested that the probability of cluster survival in the vasculature (*𝒫_C_*) is 23 to 50 times higher than that for single CTCs (*𝒫_S_*), in the next simulations, we examined the effects of changing the probability of cluster survival in the vasculature to be *𝒫_C_* = 23*𝒫_S_* = 1.15 × 10^−^ — so to take the value of the lower rather than the upper bound suggested by the authors. For this purpose, we did not allow cancer cell clusters to break up in the vasculature. Averaged over 12 simulations, the observed cluster survival was caused to be changed from 2.503 × 10^−2^ (with *𝒫* = 2.5 × 10^−2^) to 1.137 × 10^−2^. While this change had no significant effect on the number of single cells and clusters intravasating, it did reduce the average number of extravasating cancer cell clusters, as expected.

## The role of MMP-2

To investigate the role of MMP-2 in the spatiotemporal evolution of the cancer cells, we varied both the MMP-2 production rate and the MMP-2 diffusion coefficient.

Modifying the MMP-2 production rate to take values *Θ* ∈ {0, 0.1, 0.195, 0.3, 0.4} suggested that a lower MMP-2 production rate correlates to a higher overall cancer cell load after *~*22 days — for each cancer cell type individually as well as for both cell types combined. This resulted mainly from changes in cancer cell numbers on the primary grid. The corresponding plot in Figure 14 highlights this.

**Fig. 14:**
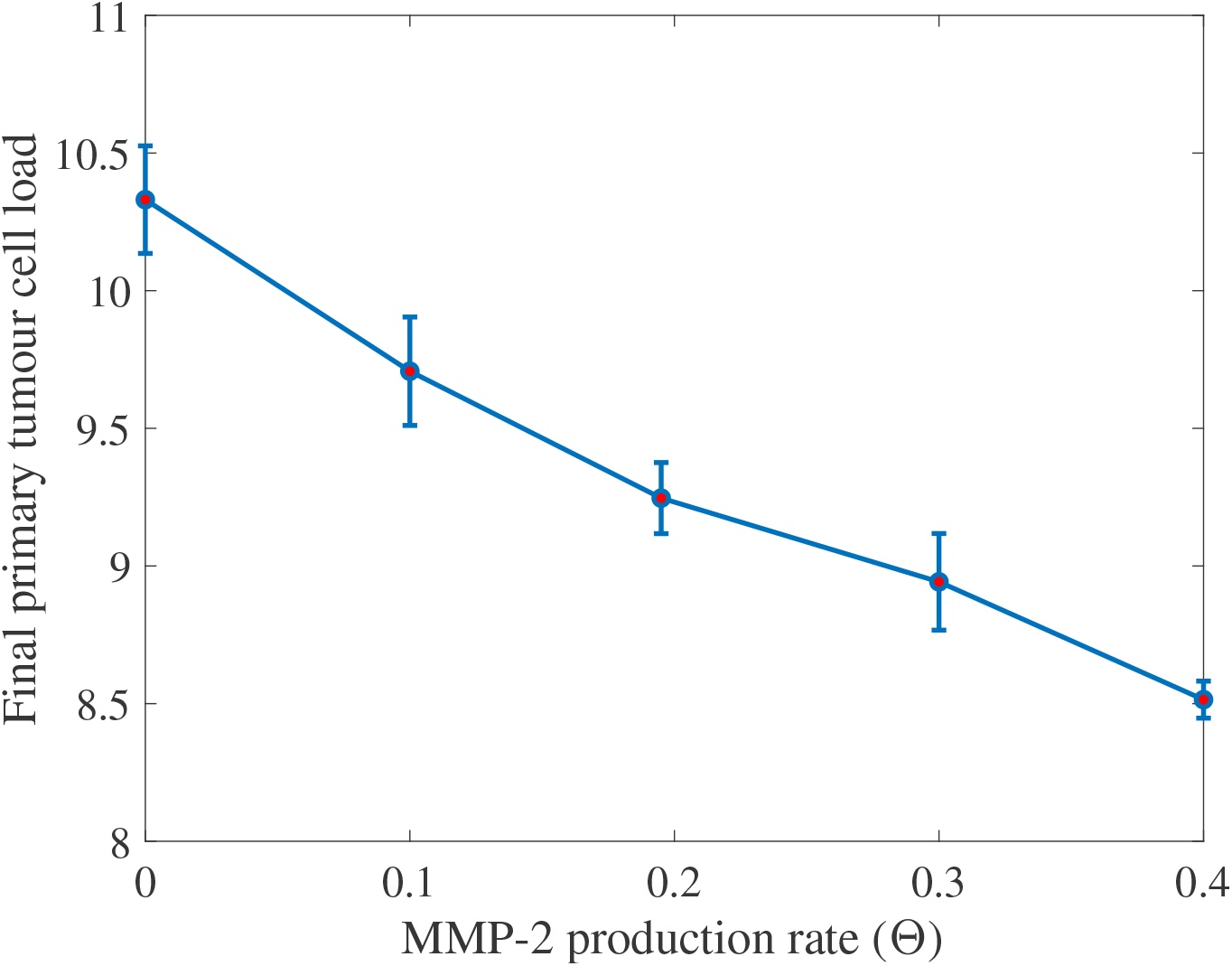
A higher MMP-2 production rate lowers the final primary tumour cancer cell load. The final primary tumour cancer cell load on the vertical axis is given in units of 10^4^ and refers to simulation results after approximately 22 days.

Increasing the MMP-2 diffusion coefficient over the range of values *D_m_* ∈ {0.1, 0.5, 1, 1.5} × 10^−3^ decreased the total cancer cell load on the primary grid after 22 days. The total number of intravasations and, coherently, the metastatic cancer cell load decreased as well. This is shown in Figure 15.

**Fig. 15:**
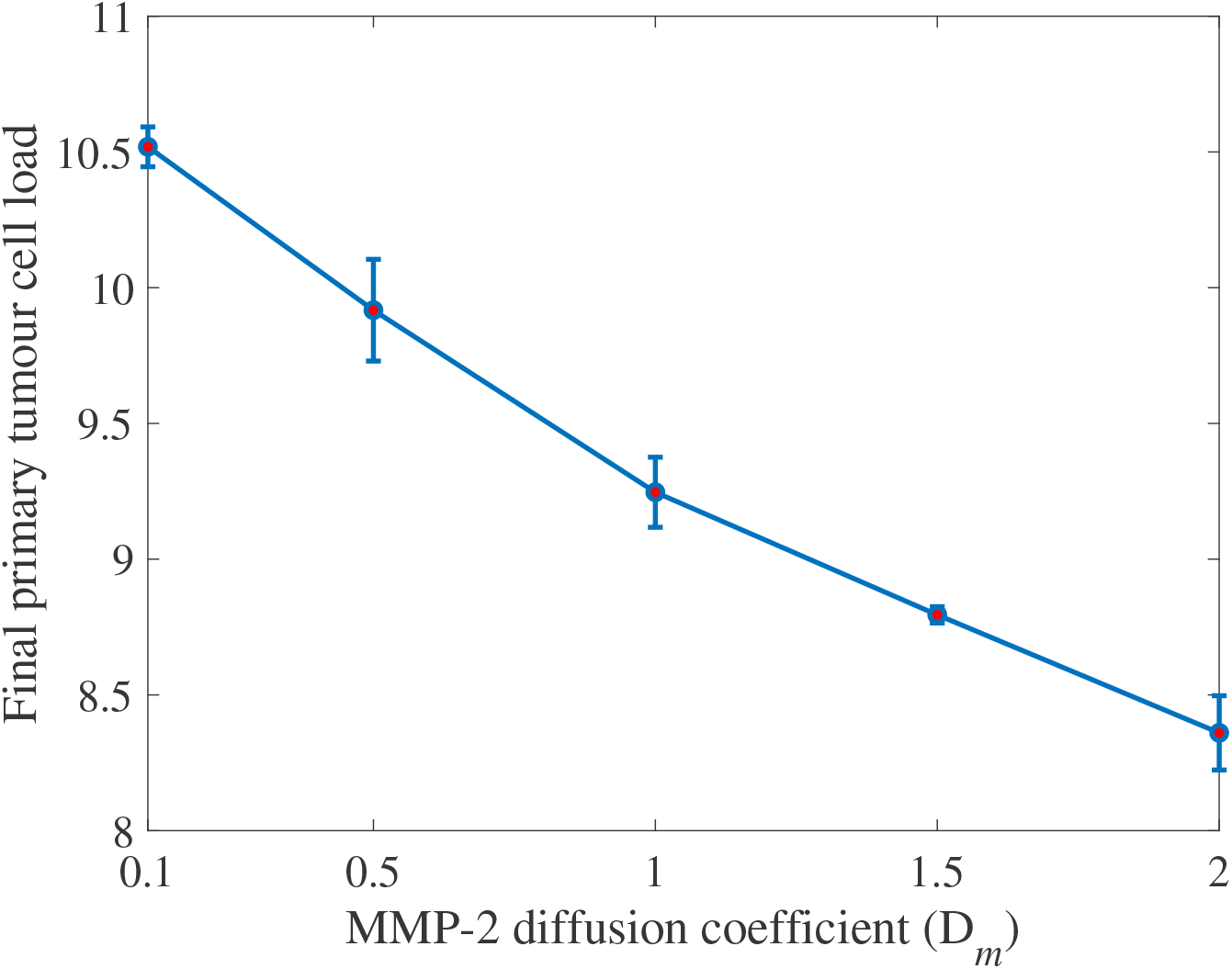
A higher MMP-2 diffusion coefficient corresponds to a lower final primary tumour cancer cell load. The MMP-2 diffusion coefficient on the horizontal axis is given in units of 10^−3^ and the final primary tumour cancer cell load on the vertical axis in units of 10^4^. The results were measured after approximately 22 days.

## The effects of MMP-2 degradation alone

We next set the MT1-MMP degradation rate to be *Γ*_1_ = 0 to examine the situation in which the diffusible MMP-2 is the only MDE in our system. We then varied the MMP-2 production rate, as we had done before when studying the effects of varying the MMP-2 production rate in the presence of MT1-MMP, to be *Θ* ∈ {0, 0.1, 0.195, 0.3, 0.4}.

Generally, we found that the total primary cancer cell load after 22 days was significantly reduced compared to simulations in which MT1-MMP was present. For instance, comparing against simulations with our baseline MMP-2 production rate of *Θ* = 0.195, the total primary cancer cell load was between 8.2% and 58.0% lower. However, invasion was still possible.

## The role of MDEs in the context of haptotaxis-dominated cancer cell movement

In all of the above simulations, we have considered diffusion-dominated cancer cell movement. We next investigated the roles of MT1-MMP and MMP-2 in cancer cell invasion in the scenario of haptotaxis-dominated cancer cell movement. For this, we changed our epithelial-like and mesenchymal-like cancer cell diffusion coefficients to be *D*_E_ = 5 × 10^−11^ and *D*_M_ = 1 × 10^−10^, respectively. Further, we focussed on cancer cell invasion on the primary grid in these simulations and hence set the number of normal and ruptured vessels to zero. *Ceteris paribus*, we then re-examined the effectiveness of invasion involving solely MT1-MMP as well as solely MMP-2 in a system with haptotaxis-dominated cancer cell movement. We first set the MT1-MMP degradation rate to be *Γ*_1_ = 0, allowing us to represent the situation in which the diffusible MMP-2 is the only MDE in our system. We then, as before, varied the MMP-2 production rate to be *Θ* ∈ {0, 0.1, 0.195, 0.3, 0.4}. As opposed to our findings when studying diffusion-dominated cancer cell movement, we observed that invasion was no longer possible for the same range of MMP-2 production rates. The final cancer cell numbers on the primary grid averaged below a ten-fold increase in cell population when compared to the original nodule of 388 cancer cells. Moreover, the final cancer cell constellation was located at the centre of the grid due to the cancer cells’ very low invasion distance.

When we increased the MT1-MMP degradation coefficient back to the baseline *Γ*_1_ = 1 but set the MMP-2 production rate to be *Θ* = 0, we found that the cancer cells did invade with an average total of 18312 cancer cells after approximately 22 days. By decreasing the MT1-MMP degradation coefficient to *Γ*_1_ = 0.5, we observed an even larger cancer cell load of 28157.

## Simulation results coincide with experimental evidence that stress importance of MT1-MMP in cancer invasion

We next ran simulations with an initial cell distribution and domain size that matched the experiments conducted by Sabeh et al (2009), who embedded HT-1080 cancer cells into 3D type I collagen gels as central nodules of diameter 1.5 × 10^−2^–2 × 10^−2^ cm. Coherently, we increased the diameter of our initial centred quasi-circular nodule to 1.5 × 10^−2^ cm and let it consist of 700 cancer cells, 40% (i.e. 280) of which were epithelial-like and 60% (i.e. 420) mesenchymal-like. Further, we decreased our domain size to be 0.1 cm × 0.1 cm to match that in the experimental conditions of Sabeh et al (2009). Figure 16 shows a snapshot of the spatio-temporal evolution of epithelial-like and mesenchymal-like cancer cells under these experimental conditions after running our simulation for 720 time steps, which corresponds to 16 days. As expected, we still observed that invasion by both epithelial-like and mesenchymal-like cancer cells was possible with both MDEs present (i.e. with *Θ* = 0.195, *Γ*_1_ = 1) or solely MT1-MMP present (i.e. with *Θ* = 0, *Γ*_1_ = 1), while invasion was not possible when solely MMP-2 was expressed (i.e. with *Θ* = 0.195, *Γ*_1_ = 0). These results are shown in the left, middle and right column of panels of Figure 16, respectively. Out of the three mechanisms, invasion under expression of MT1-MMP alone yielded the highest average invasion depth. We further observed that the switch from diffusion-dominated to haptotaxis-dominated cancer cell movement triggered more prominent finger-like protrusions in the invasive pattern of the epithelial-like cancer cells in the scenarios where either both MDEs or MT1-MMP alone were present, which are shown on the left and in the middle panel of the top row of Figure 16, respectively.

**Fig. 16:**
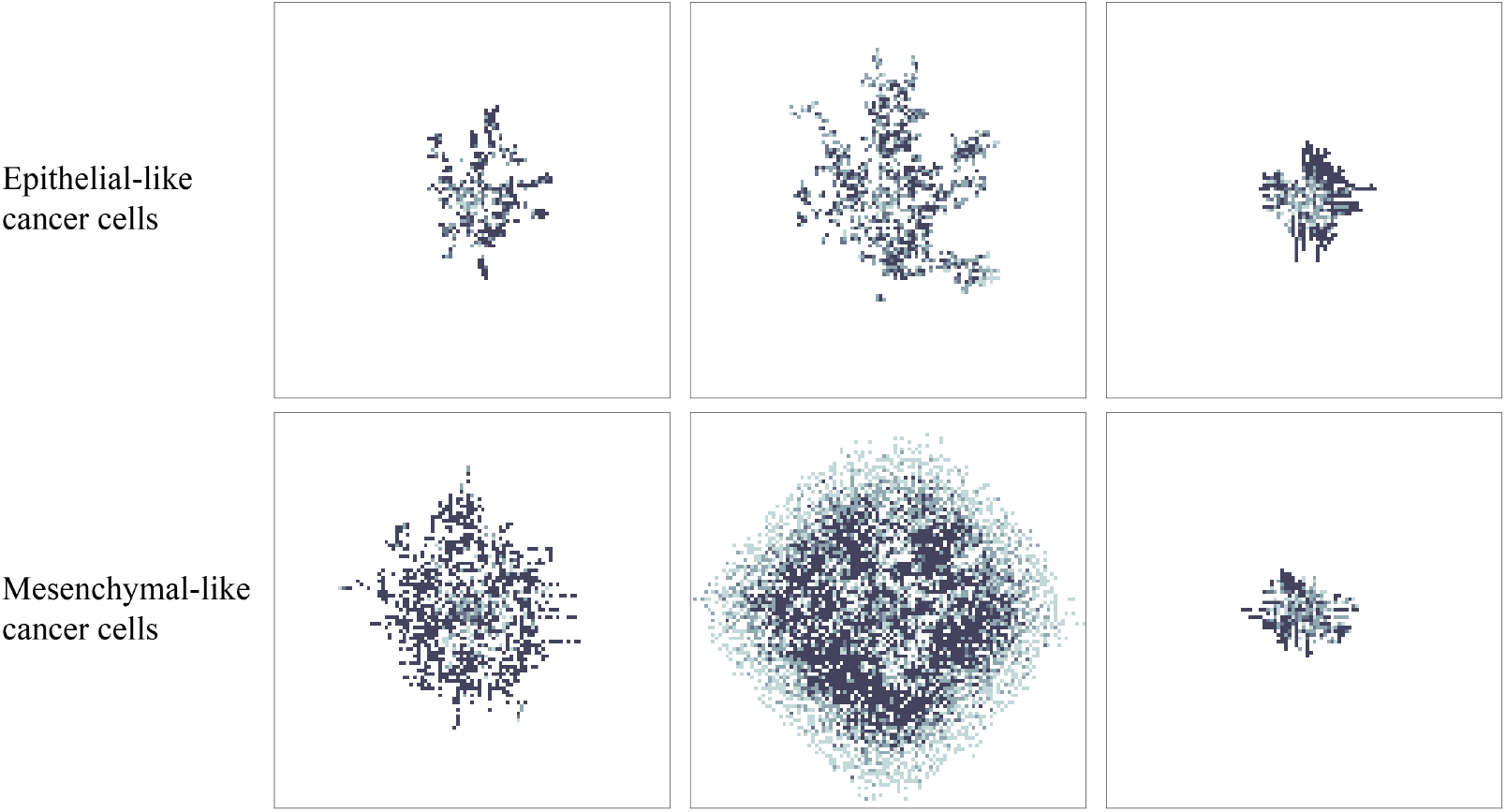
Simulation results for a heterogeneous cancer cell population subject to haptotaxis-dominated movement. To match the domain size and initial cell count of experiments by Sabeh et al (2009), we started our simulation by placing 420 mesenchymal-like cancer cells and 280 epithelial-like cancer cells in a quasi-circular region with diameter 1.5 × 10^−2^ cm at the centre of a 0.1 cm × 0.1 cm grid (initial conditions not shown). Depicted is the distribution of epithelial-like (upper panels) and mesenchymal-like (lower panels) cancer cells at time 34560Δt, corresponding to 16 days. Left to right, the invasive patterns in presence of both MDEs (Θ = 0.195, Γ_1_ = 1), in presence of MT1-MMP only (Θ = 0, Γ_1_ = 1) and in presence of MMP-2 only (Θ = 0.195, Γ_1_ = 0) are shown for both cancer cell phenotypes.

However, Sabeh et al (2009) used a *homogeneous* HT-1080 cancer cell population, and thus cells of mesenchymal origin, in their experiments to examine the role of MT1-MMP and MMP-2 in cancer invasion. In order to fully match the experimental conditions of (Sabeh et al, 2009), we next changed our initial conditions to consider a cancer cell population consisting of 700 mesenchymal-like cancer cells only, as shown in the third row of panels in Figure 17. The authors of Sabeh et al (2009) electroporated multicellular clusters of HT-1080 cancer cells of diameter 1.5 × 10^−2^–2 × 10^−2^ cm either with a control siRNA, which leaves the diffusible MMP-1 and MMP-2 as well as the non-diffusible MT1-MMP activated; or with MMP-1 and MMP-2 siRNAs, which leaves MT1-MMP alone activated but silences the diffusible MDEs MMP-1 and MMP-2; or with MT1 siRNA, which silences MT1-MMP but leaves MMP-1 and MMP-2 activated. These electroporated multicellular clusters were then embedded centrally in 3D type I collagen gels. The initial experimental setups and their respective evolution after 3 days is shown — left to right — in the two upper rows of panels in Figure 17. Since the evolution of the two diffusible MDEs, MMP-1 and MMP-2, in the experiments can be jointly accounted for by the MMP-2 equation of our model, our modelling framework can replicate the above-described experimental settings well. We did this by, again, considering a system with both MDEs (i.e. with *Θ* = 0.05, *Γ*_1_ = 1), with MT1-MMP only (i.e. with *Θ* = 0, *Γ*_1_ = 1) and with MMP-2 only (i.e. with *Θ* = 0.05, *Γ*_1_ = 0), respectively. When looking at the invasion results after 720 time steps, which corresponds to 16 days, we observed similar results as in the case of a mixed initial cancer cell population. As the first panel of the bottom row of Figure 17 shows, invasion was possible with both MDEs present. Yet, the invasion depth was slightly lower compared to the case where we allowed for MT1-MMP expression alone, which is shown in the second panel of the bottom row of the same figure. Finally, we again found that invasion was not possible in the presence of MMP-2 alone as the third panel of the bottom row of Figure 17 shows. As Figure 17 suggests, these results are qualitatively in good agreement with the experiments by Sabeh et al (2009).

**Fig. 17:**
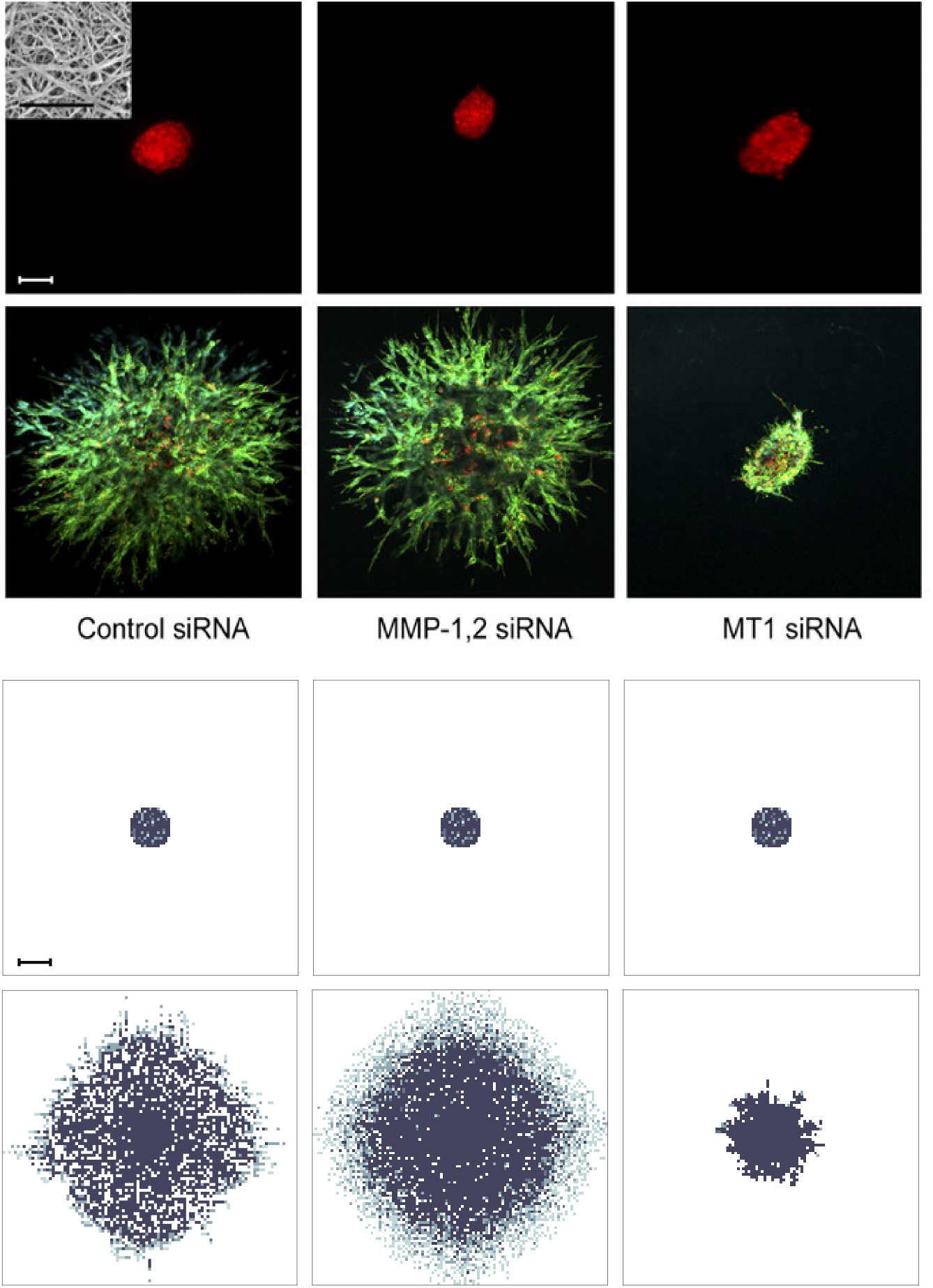
Experimental findings by Sabeh et al (2009) (black panels) compared to simulation results (white panels) The top row of panels shows the initial experimental conditions in (Sabeh et al, 2009), where HT-1080 cancer cells were embedded into 3D type I collagen gels as central nodules of diameter 1.5 × 10^−2^–2 × 10^−2^ cm. The cancer cell spheroids had previously been electroporated with a control siRNA; MMP-1 and MMP-2 siRNAs; MT1 siRNA (left to right). Their invasion after 3 days is shown in the second row of panels. To match the domain size and initial experimental conditions of Sabeh et al (2009), we started our simulation by placing 700 mesenchymal-like cancer cells in a quasi-circular region with diameter 1.5 × 10^−2^ cm at the centre of a 0.1 cm × 0.1 cm grid (third row of panels)—the bar represents a length of 1 × 10^−2^ cm. The bottom row of panels shows the distribution of the mesenchymal-like cancer cells after 16 days (i.e. at time 34560Δt), in the case where the mesenchymal-like cancer cells are subject to haptotaxis-dominated movement. Left to right, the invasive patterns in presence of both MDEs (Θ = 0.05, Γ_1_ = 1), in presence of MT1-MMP only (Θ = 0, Γ_1_ = 1), and in presence of MMP-2 only (Θ = 0.05, Γ_1_ = 0) are shown. Comparing the six panels on the bottom with those on the top, we find the simulation results to be in good qualitative agreement with the experimental results by Sabeh et al (2009).

Finally, due to our awareness that Sabeh et al (2009) used cancer cells of mesenchymal origin only in their experiments, we further sought to reproduce the results of experiments by Gaggioli et al (2007), who studied the invasion of squamous cell carcinoma (SCC) cells, which are of epithelial origin. They found that SCC cells rely on fibroblasts for their invasion, which are cells of mesenchymal-like type that are capable of matrix remodelling, like the mesenchymal-like cancer cells in our model. In the absence of fibroblasts, the authors observed that the cancer cells were unable to invade. While we showed that a mixed population of epithelial-like and mesenchymal-like cancer cells was indeed able to invade when both MDEs are present (see panels on the left of Figure 16), we ran the same simulations again with epithelial-like cancer cells only. This showed that the epithelial-like cancer cells in our model were unable to invade on their own, as observed experimentally by Gaggioli et al (2007). (Data not shown — similar to bottom right panel of Figure 17.)

## 6 Discussion and perspective

In this paper, we have presented a novel mathematical framework to model the metastatic spread of cancer in a spatially explicit manner. For this, we have used a hybrid approach (*cf*. Anderson and Chaplain (1998); Anderson et al (2000)) that captures individual cancer cell dynamics while treating abiotic factors as a continuum. We have confirmed through our computational simulations that this framework accounts for the key steps of the spatial modelling of metastatic spread.

In carrying out the computational simulations, we found that such a modelling framework provides biologically realistic outcomes and gives further insight into the mechanisms underpinning the invasion-metastasis cascade at the cellular scale. Tumour shape and metastatic distribution were predicted to appear as one would expect in a cancer patient who has not yet received treatment. In particular, we found that the mesenchymal-like cancer cells formed a ring-shaped leading front along the tumour edge, which was also seen in experiments (Nurmenniemi et al, 2009) and is often described as a leader-follower phenomenon in the literature. Nurmenniemi et al (2009) further observed an average maximum invasion depth of 5.47 × 10^−2^ cm over 14 days, when culturing HSC-3 cancer cells, a human oral squamous carcinoma cell line with high metastatic potential, on top of myoma tissue. This translates into an average maximum invasion speed of approximately 4.52 × 10^−8^ cm s^−1^. It suggests that our observed maximum invasion depth of *~*0.13 cm in 22 days and resulting average maximum invasion speed of approximately 6.77 × 10^−8^ cm s^−1^ is a realistic result, given that migration speed varies between cancer cell lines. The distribution of the cancer cell spread between the secondary sites that we considered in our model, measured via the metastatic cancer cell load and number of metastases at the respective sites, further matched the clinical data of 4181 breast cancer patients, summarised in Figure 4, which underlie our simulations. As Figure 9 indicates, the largest micrometastasis, which resulted from the earliest metastatic spread, occurred at the site of the bones. This is the most frequently observed site of metastatic spread from primary breast cancer in the data processed by the Kuhn Laboratory (2017). Overall, we observed two further successful extravasations to the site of the bones, resulting in both phenotypically homogeneous and heterogeneous secondary growth at this site. The second heaviest cancer cell load, in the form of one heterogeneous micrometastases and one consisting of epithelial-like cancer cells only, was found in the lungs. The least metastatic spread occurred to the liver with only three mesenchymal-like DTCs being observed, which arrived jointly as part of the same successfully extravasated mesenchymal-like cancer cell cluster. While, of course, stochasticity underpins the results of the metastatic spread, we again found that our results matched the clinical observations summarised in Figure 4. To our knowledge, there are currently no data available that claim to deliver an accurate estimation of the typical metastatic load from primary breast cancer to secondary sites over a specified time frame. However, we believe our result is biologically appropriate with regards to its timing, in correspondence with the conclusion reached by Obenauf and Massagué (2015) in their review of the metastatic traits that allow cancer cells to colonise various secondary sites, who suggested that CTCs and metastatic spread can be detected soon after vascularisation of the primary tumour, as in our simulations. Nonetheless, Obenauf and Massagué (2015) argue that the most limiting step of the invasion-metastasis cascade is not the dissemination through the vasculature, which we account for in our model, but the transition from infiltration of a secondary site to overt colonisation. To achieve this final step of colonisation, which is not (yet) part of our modelling framework, the cancer cells need to survive secondary site-derived detrimental signals and simultaneously exploit secondary site-derived survival signals (Obenauf and Massagué, 2015). Also, observed dormancy of tumours over extended periods may occur in the form of pre-angiogenic micrometastases that, at a later point in time, acquire the ability to become vascularized (Chambers et al, 2002). As avascular tumours can grow up to 0.1–0.2 cm via diffusion only (Folkman, 1990), all metastatic spread observed in our simulation falls into this pre-angiogenic category.

In addition to obtaining expected simulation outcomes on the cell-level, both for primary tumour growth and secondary spread, by using the baseline parameter settings in Table 1, we obtained other biologically realistic and relevant results from the simulations by varying key parameters.

Changing the initial ratio of mesenchymal-like cancer cells to epithelial-like cancer cells emphasised the importance of co-presence of the two cancer cell types for rapid invasive tumour growth. In particular, it highlighted that cancer cell invasion relies on the expression of MDEs (and MT1-MMP in particular), which are required to clear the collagen in the normal tissue of its covalent cross-links, as proposed by Sabeh et al (2009). These results further suggested that a relatively small percentage of MDE-expressing cancer cells sufficed to induce rapid cancer cell invasion.

We observed that higher numbers of mesenchymal-like cancer cells at the primary tumour location increased the number of cancer cell intravasations. Also, cancer cells from primary tumours consisting of a homogeneous epithelial-like cell population did not intravasate (unless ruptured vessels were present, in which case minimal shedding occurred). This coincides with experimental findings by Tsuji et al (2009). Results from their mouse model indicated that cancer cells originating from primary tumours of homogeneously mesenchymal-like phenotype could intravasate. The same applied to cancer cells that stemmed from tumours consisting of a combination of epithelial-like and mesenchymal-like cancer cells. On the contrary, no intravasations were observed when the primary tumour consisted of epithelial-like cancer cells only.

The fact that we, as opposed to Tsuji et al (2009), found a small amount of successfully intravasated cancer cells, even when our tumour consisted of epithelial-like cancer cells only, was to be expected. This was a result of our inclusion of ruptured vessels in the model, which we considered in accordance with the biological findings by Bockhorn et al (2007). Since these blood vessels are ruptured due to trauma or pressure applied from the expanding tumour, no MDEs are required to degrade the vessel wall and thus any cell type can intravasate through a ruptured blood vessel. We verified that the ruptured vessels were indeed the cause of this discrepancy by rerunning the simulations with an initial nodule consisting of epithelial-like cancer cells only on a primary grid that solely contained normal blood vessels.

Furthermore, we showed that our model was able to reproduce the survival probabilities of single CTCs and of CTC clusters observed in experiments by (Luzzi et al, 1998; Aceto et al, 2014).

Regarding the role of the MDEs in our model, we found that both less MMP-2 production as well as less MMP-2 diffusion caused faster cancer cell invasion and thus a higher metastatic cancer cell load after 22 days. If the MMP-2 was too diffusive or too abundant, it degraded the ECM very rapidly. The result was a ring-shaped area around the tumour edge, in which the ECM was fully degraded. This caused the influence of haptotaxis on cancer cell movement to diminish or even to cease completely. Hence, a more local or decreased degradation caused the cancer cells to invade the tissue more rapidly.

When we reduced the MMP-2 production rate to zero, we observed that the cancer cells could effectively invade in the presence of MT1-MMP only, which coincides with the experimental results by Sabeh et al (2009). On the contrary, when we set the MT1-MMP degradation rate to zero and studied the effects of MMP-2 degradation only, we found that the final total primary cancer cell load was significantly reduced compared to simulations with MT1-MMP present, which showed the same tendency as the results by Sabeh et al (2009). However, contrary to the findings in these experiments, invasion was still possible in our model when considering diffusion-dominated cancer cell movement.

To further investigate the reason for this, we reduced the diffusion coefficients of both the mesenchymal-like and the epithelial-like cancer cells, resulting in haptotaxis-dominated rather than diffusion-dominated cancer cell movement. We then studied the invasion of a mixed cancer cell population, consisting of 40% epithelial-like and 60% mesenchymal-like cancer cells, as well as of homogeneous cell populations consisting of mesenchymal-like or epithelial-like cancer cells only, under various MDE-related conditions. These conditions were the presence of MDEs as well as settings with solely MT1-MMP or solely MMP-2 present. We chose these MDE-related conditions as they correspond to those in experiments conducted by Sabeh et al (2009). We first ran simulations with a heterogeneous initial cancer cell population and then, to fully match the experimental conditions of Sabeh et al (2009), with mesenchymal-like cancer cells alone. In both cases, we found that invasion was not possible in the presence of MMP-2 alone, while invasion was possible when we considered MT1-MMP expression alone. Invasion was also possible with both MDEs present but the invasion depth was slightly decreased compared to when we considered MT1-MMP alone. In the case of a heterogeneous initial cell population, we again observed that the mesenchymal-like cancer cells formed the invading edge of the tumour by occurring most abundantly around the central cluster of epithelial-like cancer cells. Further, the epithelial-like cancer cells formed a pattern of finger-like protrusions. The simulation results observed in the case of a homogeneously mesenchymal-like cancer cell population were in qualitative agreement with the experimental results by Sabeh et al (2009). For these in vitro experiments, HT-1080 cancer cell spheroids were electroporated with a control siRNA, MMP-1 and MMP-2 siRNAs, or MT1 siRNA. These multicellular clusters were then embedded in 3D type I collagen gels as central nodules of diameter 1.5 × 10^−2^–2 × 10^−2^ cm. The invasive patterns of the HT-1080 cells under the various conditions were then studied after 3 days. Our model’s results in comparison to those of Sabeh et al (2009) are shown in Figure 17. Finally, we also matched results by Gaggioli et al (2007) confirming that in the case of an initial population of epithelial-like cancer cells only, no invasion is possible. However, the same epithelial-like population mixed with mesenchymal-like cancer cells could invade.

Since we present a first spatiotemporal modelling framework of metastatic spread, we have focussed on capturing the main steps involved in the invasion-metastasis cascade. In future work, we aim to include more biological detail.

Adding a third spatial dimension to our problem would be one natural extension. We have not prioritised this at this early stage of the development of our invasion-metastasis cascade modelling framework as we believe that this would not change the overall characteristics of the model’s qualitative insights.

Biomechanical properties are also not accounted for in our first modelling framework but are evidently important, especially for processes such as intravasation, travel through the vasculature and extravasation. For this reason, we are planning to couple our modelling framework to a biomechanical haemodynamics model.

Breast cancer cell intravasation and dissemination occurs through a *tumour microenvironment of metastasis* (TMEM)-mediated mechanism. TMEMs are microanatomical structures consisting of three different cell types that are in direct physical contact with one another (Karagiannis et al, 2017). Since we model individual cell dynamics, our framework can easily be adapted to model TMEM involvement in metastatic spread by including an additional cancer cell type and calibrating its phenotype.

Often solitary DTCs and micrometastases are non-proliferating, or *dormant*, even years after primary tumour diagnosis (Chambers et al, 2002; Pantel and Speicher, 2016). Others may not survive due to activation of the immune system at the secondary sites. We could include cancer cell dormancy and immune system activation at secondary sites more explicitly. In this regard, we could extend our framework to include site-specific differentiation of the tissue at the metastatic sites in the body. Further, the tumour microenvironment at the metastatic sites could be modelled as a landscape that evolves over time to capture the process of pre-metastatic niche formation that has been observed both in mouse models and clinical studies (McAllister and Weinberg, 2014). It could thus be studied in more detail how these changes in the dormant cancer cells’ microenvironment over time affect growth activation of previously latent micrometastases, which was suggested, amongst others, by McAllister and Weinberg (2014).

To expand our model to capture metastatic growth beyond the current avascular stage and to thus include the development of larger, vessel-growth activating (macro-)metastases, it would also be useful to include tumour-induced angiogenesis into our modelling framework. This could be achieved, for example, by accounting for tumour-secreted, diffusive TAFs that trigger angiogenesis both at the primary and the secondary sites of our model, in a similar manner to Anderson and Chaplain (1998). Incorporating tumour-induced angiogenesis in our framework would also allow us to include reseeding in a more realistic manner. In this paper, we have investigated the classic, unidirectional view of metastatic progression. However, it is hypothesised that *self-seeding* from primary to primary tumour, *primary reseeding* from a metastatic site back to the primary tumour, and *metastatic reseeding*, where metastases form out of existing metastases, also play a role in metastatic spread. These processes could easily be included in this modelling framework, in particular in the context of allowing for colonisation by considering tumour-induced angiogenesis at the secondary sites. Of course, we could also replace the currently static initial vessel distribution in a natural way by including vascular growth resulting from tumour-induced angiogenesis instead. Moreover, similarly to Powathil et al (2012), we could include the transportation of oxygen to the tumour through newly formed vessels as well as hypoxia-induced quiescence of cancer cells.

In the context of micrometastatic dormancy — and in general — mutations of the cancer cells could also easily be included in the model. In particular, modelling mutations that enable EMT and the reverse process, MET, to represent permanent and transient transitions between an epithelial and mesenchymal phenotype as observed in the invasion-metastasis cascade (Celià-Terrassa et al, 2012) could deliver valuable insight. Further, we could account for the fact that EMT has been shown to predominantly occur at the tumour edges (Tsuji et al, 2009) since our modelling technique is spatially explicit. Also, it has been suggested that mesenchymal-like and epithelial-like phenotypes occur on a spectrum rather than as two discrete groups (Campbell, 2018). Hence, it would be beneficial to present the phenotype of cancer cells as a continuum.

For ethical reasons, modern-day data concerning metastatic spread, such as those by Kuhn Laboratory (2017), stem from studies in which the primary tumours were removed prior to the observations of when the tumour would reoccur or present detectable metastases. We could easily modify our model to include both successful resections of the primary tumour and/or of metastases as well as tumour resections that accidentally leave a small residue of cancer cells. Finally, we could model resections at various times to examine the effects of delayed surgical interventions on the disease outcome.

## Acknowledgements

LCF would like to thank Jack Nichol (Abertay University) for his helpful C++-specific advice.

## Appendices

### A Non-dimensionalisation

The following dimensional continuum system of PDEs, along with zero-flux boundary conditions, describes the spatio-temporal evolution of the epithelial-like cancer cell density *c*_E_ (*t, x, y*), of the mesenchymal-like cancer cell density *c*_M_ (*t, x, y*), of the ECM density *w*(*t, x, y*), and of the MMP-2 concentration *m*(*t, x, y*):

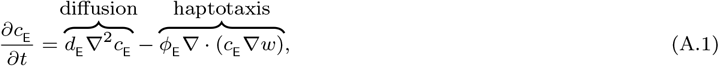

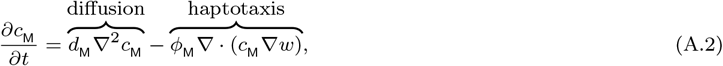

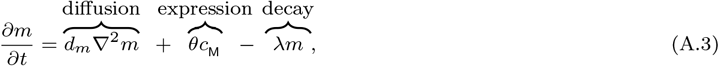

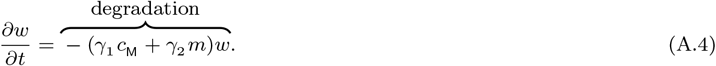

We non-dimensionalise the system in coherence with Anderson et al (2000). So we choose to rescale distance with a length scale *L* = 0.2 cm, which represents an appropriate choice for the maximum invasion distance of cancer cells at an early stage of cancer growth, and rescale time by 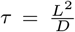, where *D* = 10^−6^ cm^2^s^−1^ is a reference chemical diffusion coefficient, such that *τ* = 4 × 104 s, which gives approximately 11 h. We non-dimensionalise our model by setting 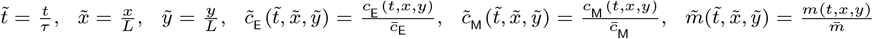 and 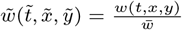 where *c*_E_, *c*_M_, *m* and *w* are appropriate reference parameters. When substituting these into the system of PDEs (A.1)–(A.4) and dropping the tildes for better readability, we obtain

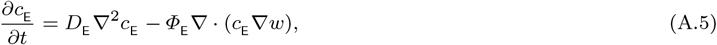

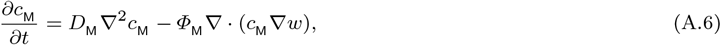

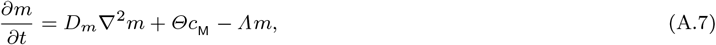

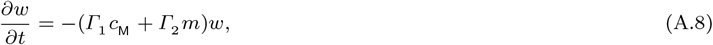

where 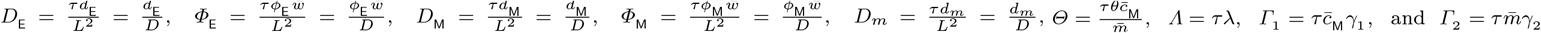

To choose biologically realistic parameter values for our model, we consult biological publications on the topic (Stokes et al, 1990; Bray, 1992; Luzzi et al, 1998; Meng et al, 2004; Milo et al, 2009; Collier et al, 2011; Vajtai, 2013; Aceto et al, 2014; Kuhn Laboratory, 2017) as well as comparable PDE models (Anderson et al, 2000; Deakin and Chaplain, 2013). An overview of the parameter values used together with their mathematical and experimental origin can be found in Table 1.

### B Hybrid approach outline

We fix a time step *Δt* and set *t_n_* = *nΔt*. We discretise each of the *G* + 1 square domains using a uniform mesh with step size *Δx* = *Δy* = 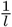. We set *x_i_* = *iΔx* and *y_j_* = *jΔy*, where *i, j ∈* [0, *l*] *⊂* ℕ_0_. We continue by approximating the MMP-2 concentration *m*(*t, x, y*) and the ECM density *w*(*t, x, y*) by discrete values 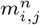 and 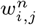, respectively, and denote the number of epithelial- and mesenchymal-like cancer cells on grid point (*x_i_, y_j_*) at time *t_n_* by 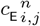 and 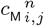, respectively.

The spatio-temporal evolution of MMP-2 concentration and ECM density is then governed by the explicit five-point central difference discretisation of equations (3) and (4), respectively:

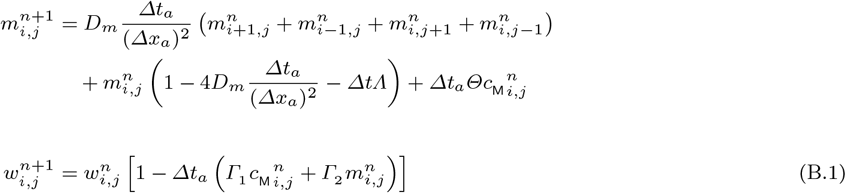

along with zero-flux boundary conditions. Note that the discretisation time step *Δt* and space steps *Δx* and *Δy* were chosen to represent the physical properties of cancer cell size and remain fixed. The abiotic time step *Δt_a_* and the abiotic space steps *Δx_a_* and *Δy_a_* can be chosen freely for a more accurate discretisation of the PDEs in (B.1), as long as *Δt*, *Δx* and *Δy* are integer multiples of *Δt_a_*, *Δx_a_* and *Δy_a_*, respectively.

We discretise equations (1) and (2) using, again, a five-point central difference scheme. This yields

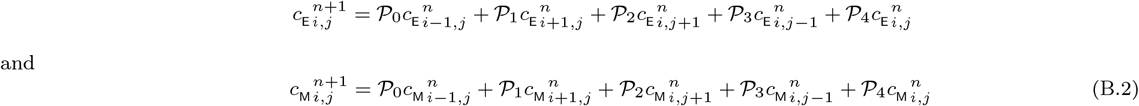

for epithelial-like and mesenchymal-like cancer cells, respectively, together with zero-flux boundary conditions.

We then extract the coefficients of the resulting probabilities for cancer cells to move left, right, up and down to determine the cancer cells’ movement. We obtain

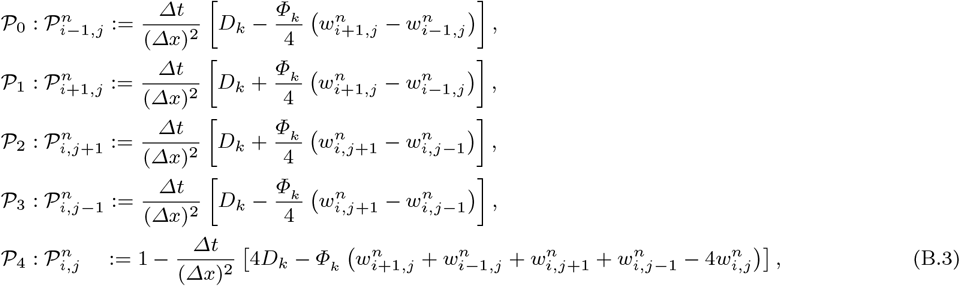

where *k* = E, M, with the coefficients *𝒫*_0_, *𝒫*_1_, *𝒫*_2_, *𝒫*_3_ corresponding to the probabilities that, during the next time step, a cancer cell at grid point (*x_i_, y_j_*) moves left, right, up and down, respectively. While the above probabilities *𝒫*_0_–*𝒫*_4_ let us derive equations (1) and (2), we will choose for *P*_4_ to become

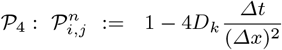

in order to make *𝒫*_0_–*𝒫*_4_ real probabilities such that

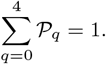

*𝒫*_4_ is the probability that a cancer cell remains at grid point (*x_i_, y_j_*) during the next time step. So the cancer cells move both by diffusion, and, as soon as there is a non-zero ECM density gradient, also by haptotactic movement towards the higher ECM density.

At any time step *n ≥* 0, we realise the individual-based cell movement from grid point to grid point using the following algorithm, following the work of Burgess et al (2016, 2017):

1. On every grid we identify 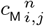 on each grid point (*x_i_, y_j_*) by counting the number of mesenchymal-like cancer cells and thus the MMP-2 concentration and ECM density by calculating the numerical solutions defined by completing equations (B.1) with zero-flux boundary conditions and suitable initial conditions, which we defined above.
2. For each grid point (*x_i_, y_j_*) on every grid, we evaluate the movement probabilities to a neighbouring grid point for cancer cells on this grid point by substituting the local ECM densities into equations (B.3).
3. Five intervals are then defined based on the movement probabilities in equations (B.3), at each grid point (*x_i_, y_j_*):

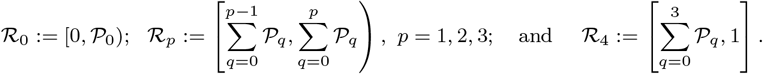
4. At each grid point (*x_i_, y_j_*) of every grid, we generate a random number *z ∈* [0, 1] for every cancer cell on that grid point. Depending on which of the above intervals 𝓡_0_ to 𝓡_4_ the value of *z* falls into, the corresponding cancer cell will move left (*z ∈ 𝓡*_0_), move right (*z ∈ 𝓡*_1_), move up (*z ∈ 𝓡*_2_), move down (*z ∈ 𝓡*_3_), or remain on its current grid point (*z ∈ 𝓡*_4_).
5. If a cancer cell would have been placed outside the grid limits by Step 4, it remains in its grid position in compliance with our zero-flux boundary conditions. The same applies if a cancer cell would have moved to a grid point already filled with the maximum carrying capacity of *𝒬* cells.

### C Figures

**Fig. 18:**
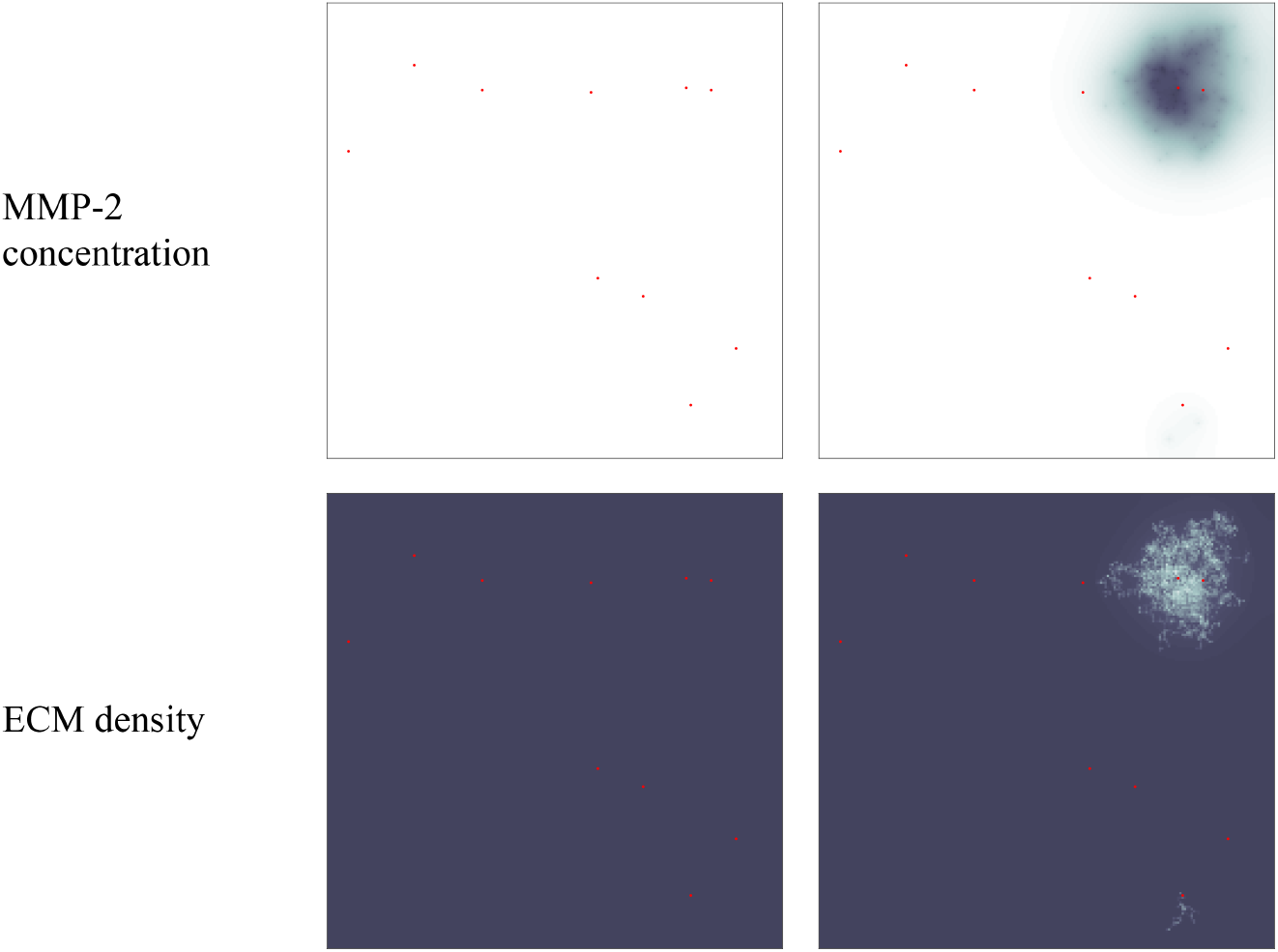
Simulation results on secondary grid representing the bones. Distribution of MMP-2 concentration (top panels) and ECM density (bottom panels) at the secondary site of the bones is shown at times 24000Δt (left) and 48000Δt (right). This corresponds to ~11 days and ~22 days, respectively. The MMP-2 concentration ranges from 0 (white) to 3.4737 × 10^−2^ (black) and the ECM density from 0.17559 (light grey) to 1 (black). The red grid points represent blood vessels, through which cancer cells can extravasate. Figure 9 shows the corresponding plots of the cancer cell distributions.

**Fig. 19:**
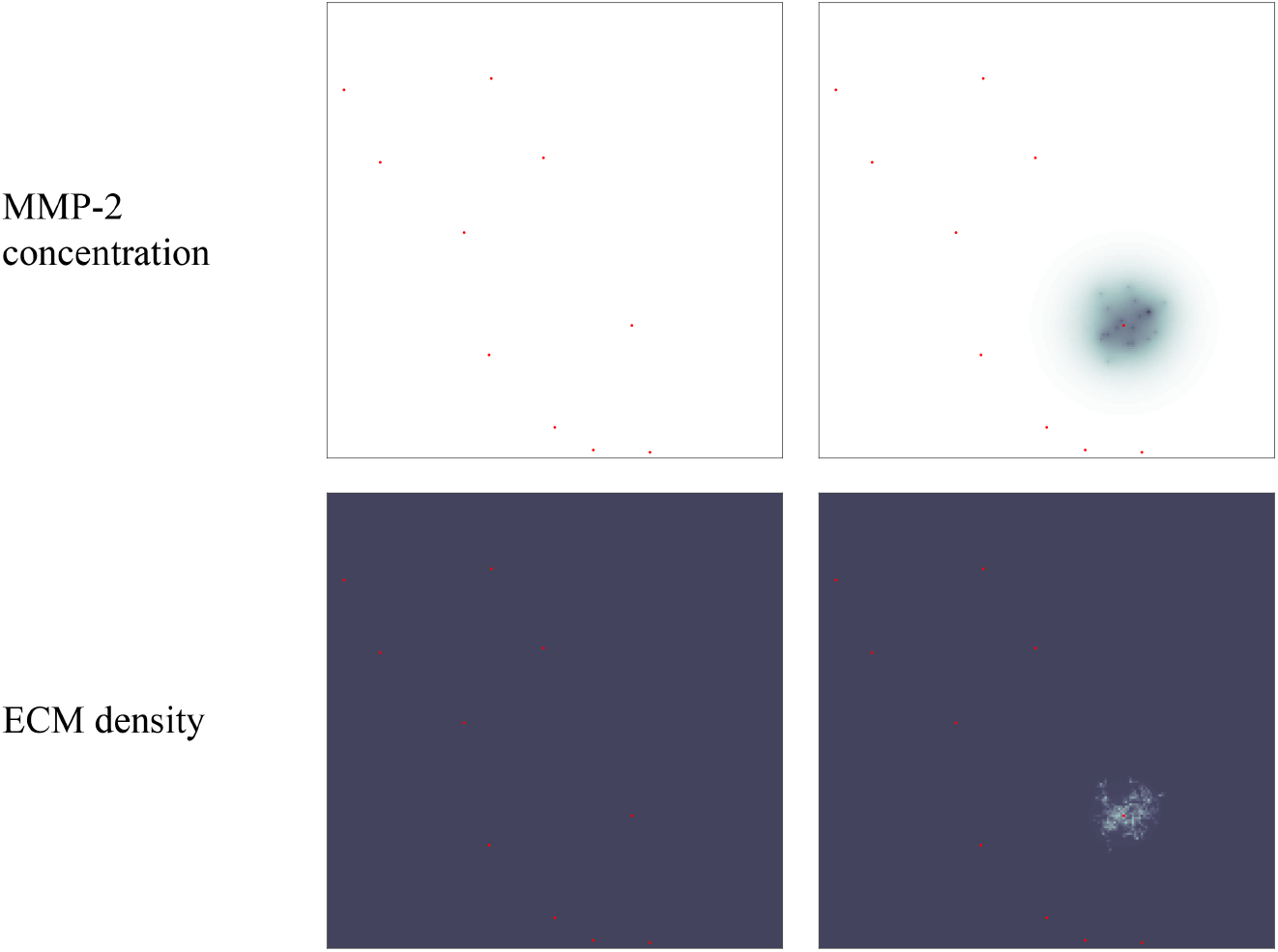
Simulation results on secondary grid representing the lungs. Distribution of MMP-2 concentration (upper panels) and ECM density (lower panels) at the secondary site of the lungs is shown at times 24000Δt (left) and 48000Δt (right). This corresponds to ~11 days and ~22 days, respectively. The MMP-2 concentration ranges from 0 (white) to 1.5876 × 10^−2^ (black) and the ECM density from 0.41137 (light grey) to 1 (black). The red grid points represent blood vessels, through which cancer cells can extravasate. Figure 10 shows the corresponding plots of the cancer cell distributions.

**Fig. 20:**
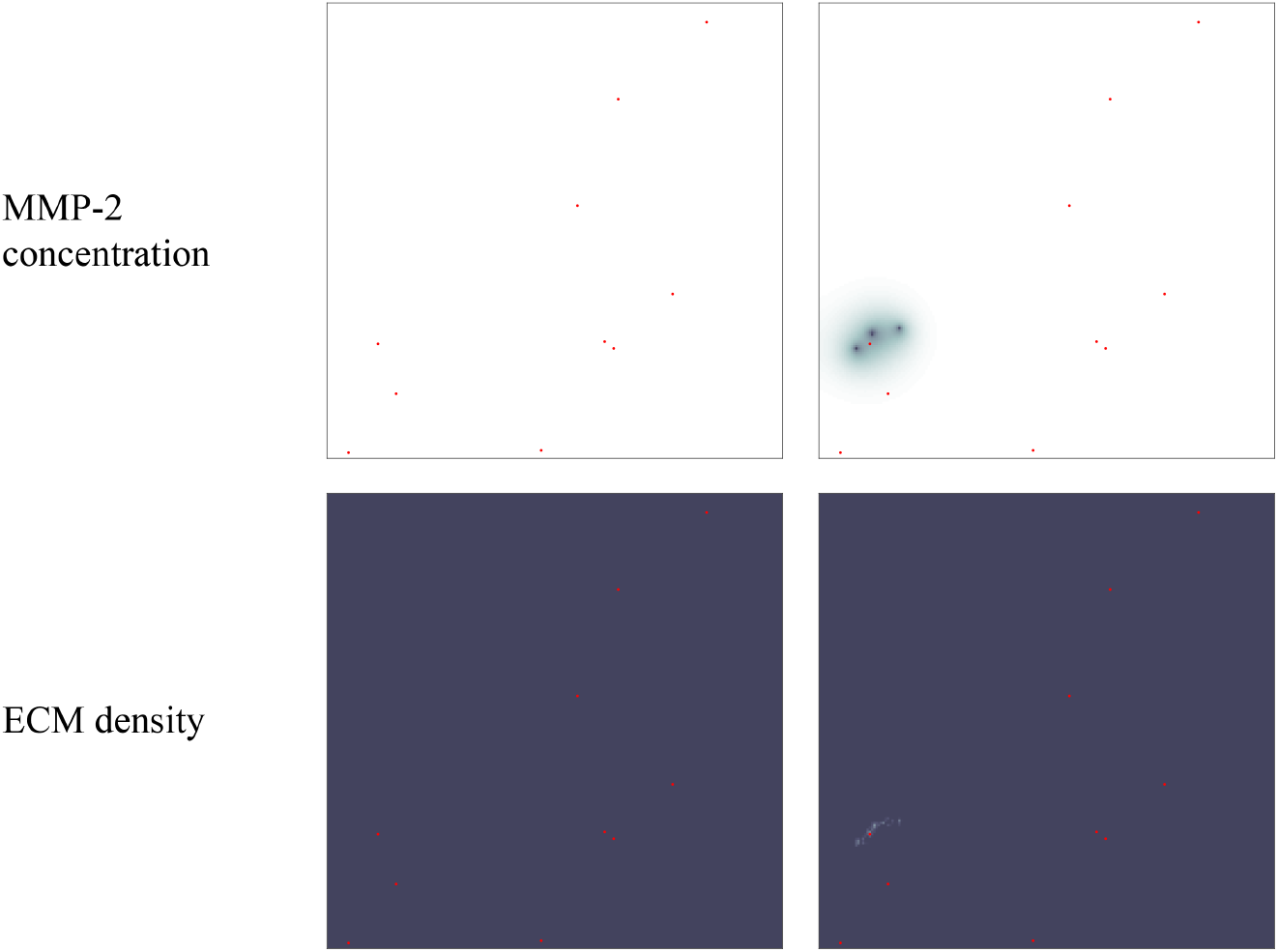
Simulation results on secondary grid representing the liver. Distribution of MMP-2 concentration (top panels) and ECM density (bottom panels) at the secondary site of the liver is shown at times 24000Δt (left) and 48000Δt (right). This corresponds to ~11 days and ~22 days, respectively. The MMP-2 concentration ranges from 0 (white) to 3.7129 × 10^−3^ (black) and the ECM density from 0.58015 (light grey) to 1 (black). The red grid points represent blood vessels, through which cancer cells can extravasate. Figure 11 shows the corresponding plots of the cancer cell distributions.

## Footnotes

1 The notation (*x_i_, y_j_*) *∈ Ω*_P_ is a result of the discretisation of the grids in our model, as described in detail in Appendix B.

